# Monocyte-derived macrophages aggravate pulmonary vasculitis via cGAS/STING/IFN-mediated nucleic acid sensing

**DOI:** 10.1101/2022.05.30.493983

**Authors:** Nina Kessler, Susanne F. Viehmann, Calvin Krollmann, Karola Mai, Katharina Maria Kirschner, Hella Luksch, Prasanti Kotagiri, Alexander M.C. Böhner, Dennis Huugen, Carina C. de Oliveira Mann, Simon Otten, Stefanie A. I. Weiss, Thomas Zillinger, Kristiyana Dobrikova, Dieter E. Jenne, Andrea Ablasser, Eva Bartok, Gunther Hartmann, Karl-Peter Hopfner, Paul A. Lyons, Peter Boor, Angela Rösen-Wolff, Lino Teichmann, Peter Heeringa, Christian Kurts, Natalio Garbi

## Abstract

Autoimmune vasculitis is a group of life-threatening diseases, whose underlying pathogenic mechanisms are incompletely understood, hampering development of targeted therapies. Here, we demonstrate that patients suffering from anti-neutrophil cytoplasmic antibodies (ANCA)-associated vasculitis (AAV) showed increased activity of the DNA sensor cGAS and enhanced IFN-I signature. To identify potential therapeutic targets, we developed a mouse model for pulmonary AAV that mimics severe disease in patients. Immunogenic DNA accumulated during disease onset, triggering cGAS/STING/IRF3-dependent IFN-I release that promoted endothelial damage, pulmonary hemorrhages, and lung dysfunction. Macrophage subsets played dichotomic roles in disease. While recruited monocyte-derived macrophages were major disease drivers by producing most IFN-β, resident alveolar macrophages contributed to tissue homeostasis by clearing red blood cells and limiting infiltration of IFN-β-producing macrophages. Moreover, pharmacological inhibition of STING, IFNAR-I or its downstream JAK/STAT signaling reduced disease severity and accelerated recovery. Our study unveils the importance of STING/IFN-I axis in promoting pulmonary AAV progression and identifies cellular and molecular targets to ameliorate disease outcome.

**Graphical abstract:** 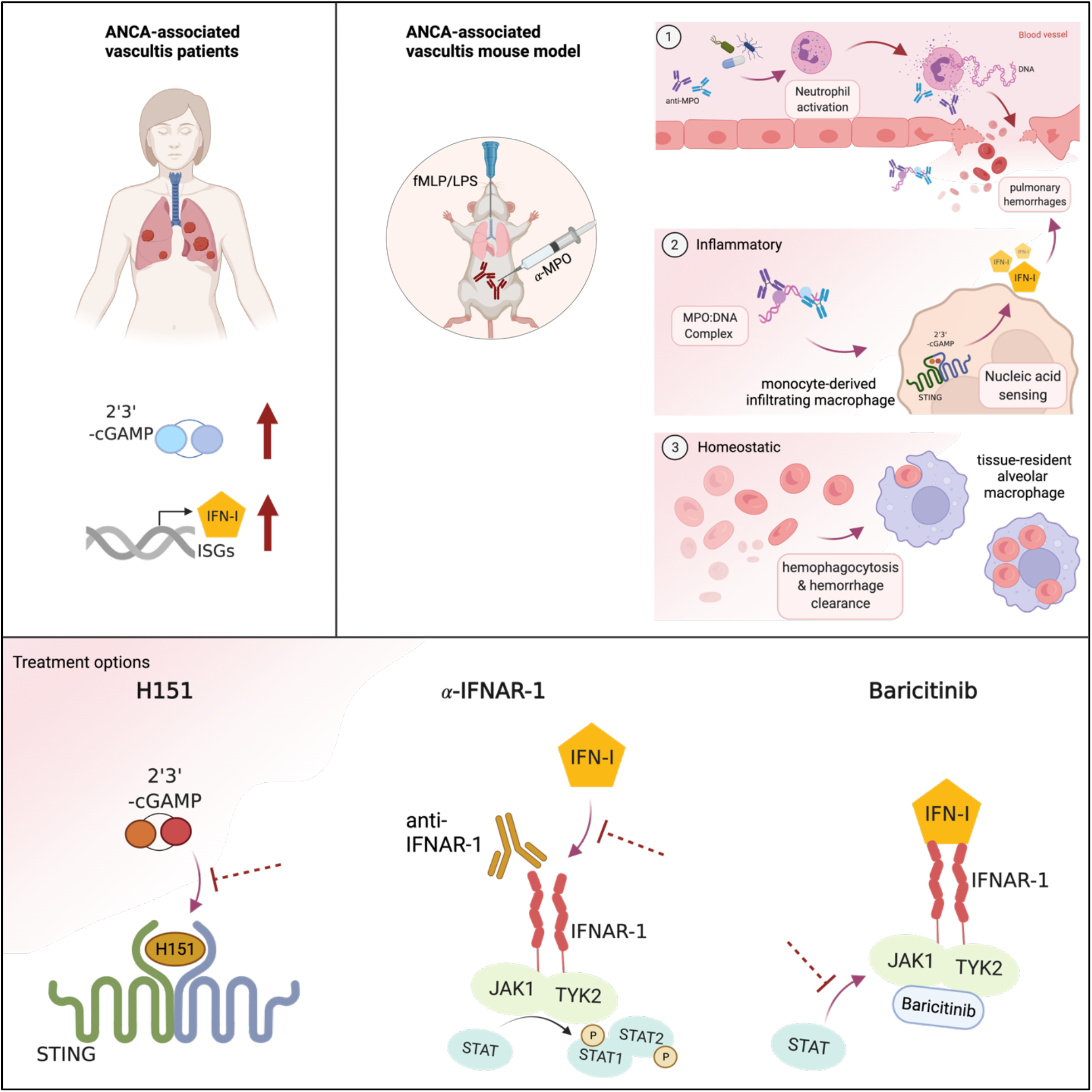

**Summary:** Kessler et al. identify aberrant DNA recognition by cGAS/STING axis and IFN-I production by inflammatory macrophages as a major driver of severe ANCA-associated vasculitis (AAV). Pharmacological interventions blocking this pathway ameliorate disease and accelerate recovery, identifying potential targets for therapeutic intervention in patients.

## Introduction

Recognition of cytosolic DNA by the cGAS/STING pathway is a key process in anti-viral immunity resulting in the release of IFN-I, TNF-α and IL-6 (Barrat, Elkon, and Fitzgerald 2016; Motwani, Pesiridis, and Fitzgerald 2019; Gao et al. 2013; Ishikawa and Barber 2008; Wu et al. 2013). Since these cytokines have an important pro-inflammatory role in different autoimmune diseases, it is conceivable that DNA released from stressed cells may activate nucleic acid sensing pathways and worsen disease progression. Aberrant DNA release is indeed characteristic of several autoimmune diseases including anti-neutrophilic cytoplasmic antibodies (ANCA)-associated vasculitis (AAV) (Kessenbrock et al. 2009; Sangaletti et al. 2012; Surmiak et al. 2015), thus posing the question whether the cGAS/STING pathway is promoting AAV progression.

AAV encompasses various diseases characterized by high levels of pathogenic ANCAs, mostly directed against myeloperoxidase (MPO) or proteinase 3 (PR3), accumulation of extracellular DNA at the sites of injury and necrotizing inflammation of small and intermediate vessels (Jennette and Falk 1997; Kessenbrock et al. 2009; O’Sullivan et al. 2015; Kitching et al. 2020). AAV is a systemic disease that often manifests in the aftermath of bacterial infections (Timmeren, Heeringa, and Kallenberg 2014). Because of the scarce knowledge on underlying pathological mechanisms, current therapies are based on broad immunosuppression, which have significantly lowered mortality in AAV patients (Rich and Brown 2012) at the expense of considerable side effects of long-term corticosteroid administration. For example, infections have become the main primary cause of death within the first year of diagnosis (47%), whereas active vasculitis itself accounts for a lower fraction of mortality (19%). The high rate of infection-related mortality together with other severe side effects (Flossmann et al. 2011; Little et al. 2010) warrant for the identification of disease pathways that can be the basis for development of novel, selective therapies.

Central to this question is the availability of robust pre-clinical models to identify key molecular and cellular mechanisms driving AAV progression (Jennette et al. 2013). Existing mouse models based on active or passive immunization against MPO have been instrumental in uncovering the pathogenic role of anti-MPO Ig (Xiao et al. 2002; Schreiber et al. 2006; Little et al. 2009) and the C5a-C5aR axis (Jayne et al. 2017; Xiao et al. 2014) as well as defining the contribution of neutrophils (Xiao et al. 2005), monocytes (Rousselle, Kettritz, and Schreiber 2017; Rousselle et al. 2022), and T cells (Gan et al. 2010). However, the high variability in disease incidence and severity inherent to those animal models considerably limits their impact on identifying novel therapeutic targets (Gan et al. 2015; Shochet, Holdsworth, and Kitching 2020). In addition, although hemorrhages in the lung are amongst the most life-threatening features of small- and medium-vessel AAV (Jennette and Falk 1997), none of the available mouse models is associated with significant pulmonary involvement. The establishment of a reproducible and clinically relevant mouse model for severe autoimmune vasculitis is thus necessary for the identification of disease mechanisms that can be therapeutically targeted.

Here we report in two independent patient cohorts that active AAV is associated with enhanced cGAS activity and an IFN-I signature in blood PBMCs, suggesting an active pathway of DNA sensing mediated by cGAS/STING/IFN-I. To investigate this, we established an ANCA-dependent mouse model of severe pulmonary hemorrhages resembling clinical features of life-threatening AAV. Using this AAV animal model, we demonstrate a role of cGAS/STING/IRF3 pathway in disease pathogenesis by promoting IFN-I release and endothelial cell damage. In addition, we identify a functional dichotomy between lung macrophages whereby recruited monocyte-derived macrophages promoted disease progression by producing most IFN-I, while resident alveolar macrophages had homeostatic function by clearing hemorrhages. Therapeutic intervention to inhibit STING activation, IFN-I recognition, or JAK1/2 signaling improved disease outcome. Thus, our results identify the cGAS/STING/IFN-I axis of nucleic acid recognition as a potential therapeutic target to ameliorate AAV.

## Results

### Increased nucleic acid sensing and IFN-I signature in patients with ANCA-vasculitis

Patients with anti-MPO-associated vasculitis have increased numbers of netting neutrophils at the sites of injury that release MPO-decorated DNA (Kessenbrock et al. 2009). However, the contribution of DNA sensing in disease pathogenesis is unknown. Here, we found increased cGAMP concentration in the serum of patients undergoing active ANCA-associated small-vessel vasculitis compared to healthy controls (Fig. 1 A and Table 1), suggesting ongoing DNA recognition by the cGAS/STING pathway. Further supporting this, we demonstrate an increased IFN-I signature during active disease in an independent patient cohort of patients with ANCA-associated vasculitis with similar clinical characteristics (Fig. 1 B and C, and Table 2). Hallmark IFN-I-induced genes including *Mx2*, *Oasl*, *Ifi27/35*, and *Arg1* amongst others were upregulated in patients undergoing active vasculitis (Fig. 1 C and D). These results suggest a potential role of cGAS/STING-dependent DNA recognition in disease progression of patients with active MPO-associated small-vessel vasculitis.

**Figure 1.**
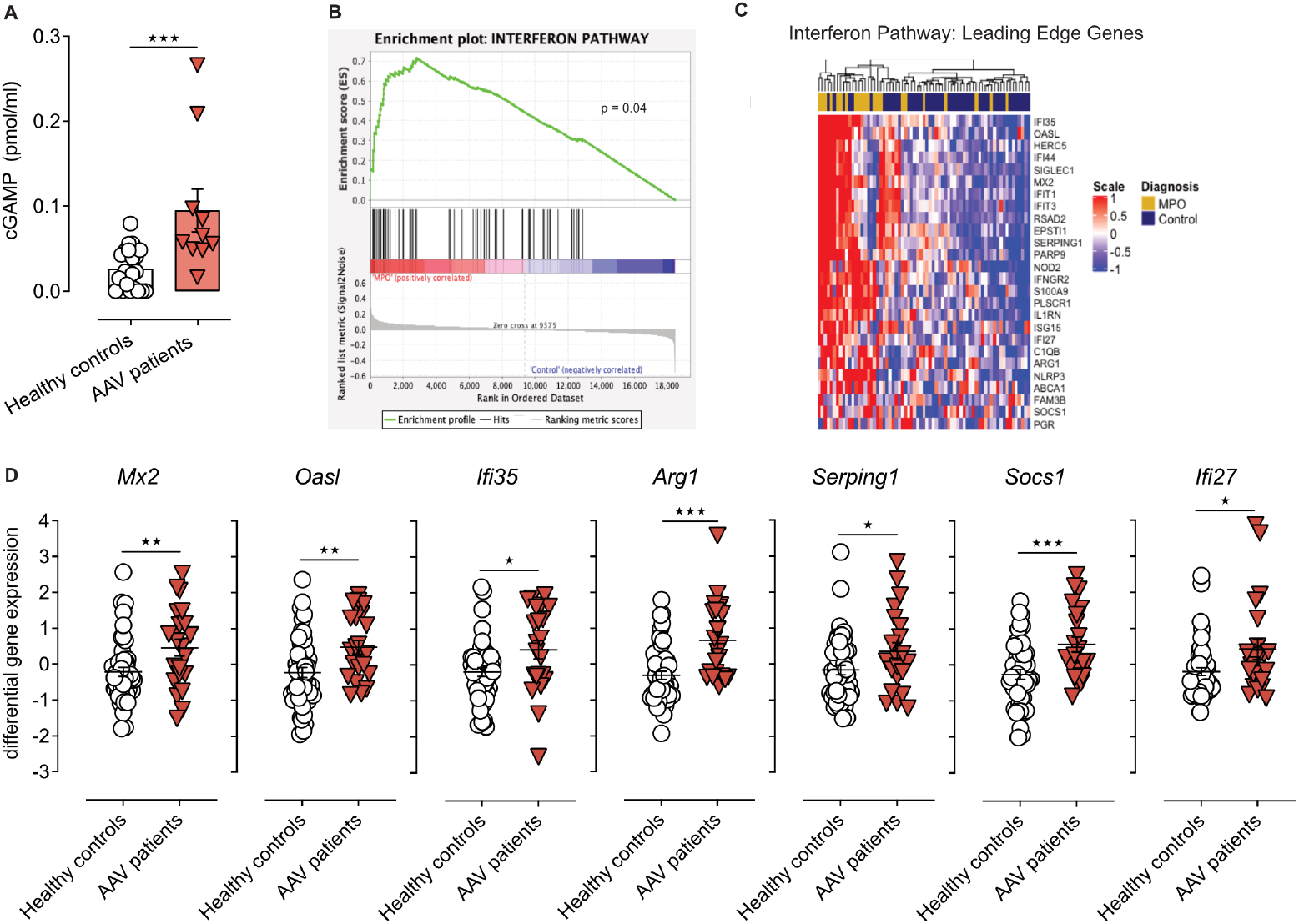
Patients with ANCA-associated vasculitis show increased cGAS activation and IFN-I signature. **(A)** cGAMP concentration in PBMCs isolated from healthy donors or patients with active anti-MPO-associated vasculitis (patient cohort I -Table 1). Each dot is an individual donor (n = 22 for healthy, n = 10 for MPO-vasculitis donors). Columns represent mean ± SEM. **(B)** Gene set enrichment analysis (GSEA) in patients undergoing active anti-MPO-associated vasculitis (n = 21, patient cohort II -Table 2). **(C)** Heatmap of IFN-I pathway leading-edge gene expression levels in patients with active anti-MPO-associated vasculitis (MPO, n = 21) or in remission (Control, n = 47) (patient cohort II). **(D)** Expression levels for selected ISGs from analysis in (C). *, P<0.05; **, P<0.01; ***, P<0.001. cGAMP, 2’3’-cyclic GMP-AMP; AAV, anti-neutrophil cytosolic antibodies-associated vasculitis; MPO, myeloperoxidase.

### Combination of bacterial ligands and anti-MPO antibodies provoke severe lung hemorrhages and dysfunction in a novel mouse model

To investigate pathophysiological mechanisms driving small vessel autoimmune vasculitis, we aimed at establishing a novel mouse model of severe ANCA-induced vasculitis in the lung, which is the most critical disease manifestations in patients (Jennette and Falk 1997) (Fig. 2 A).

**Figure 2.**
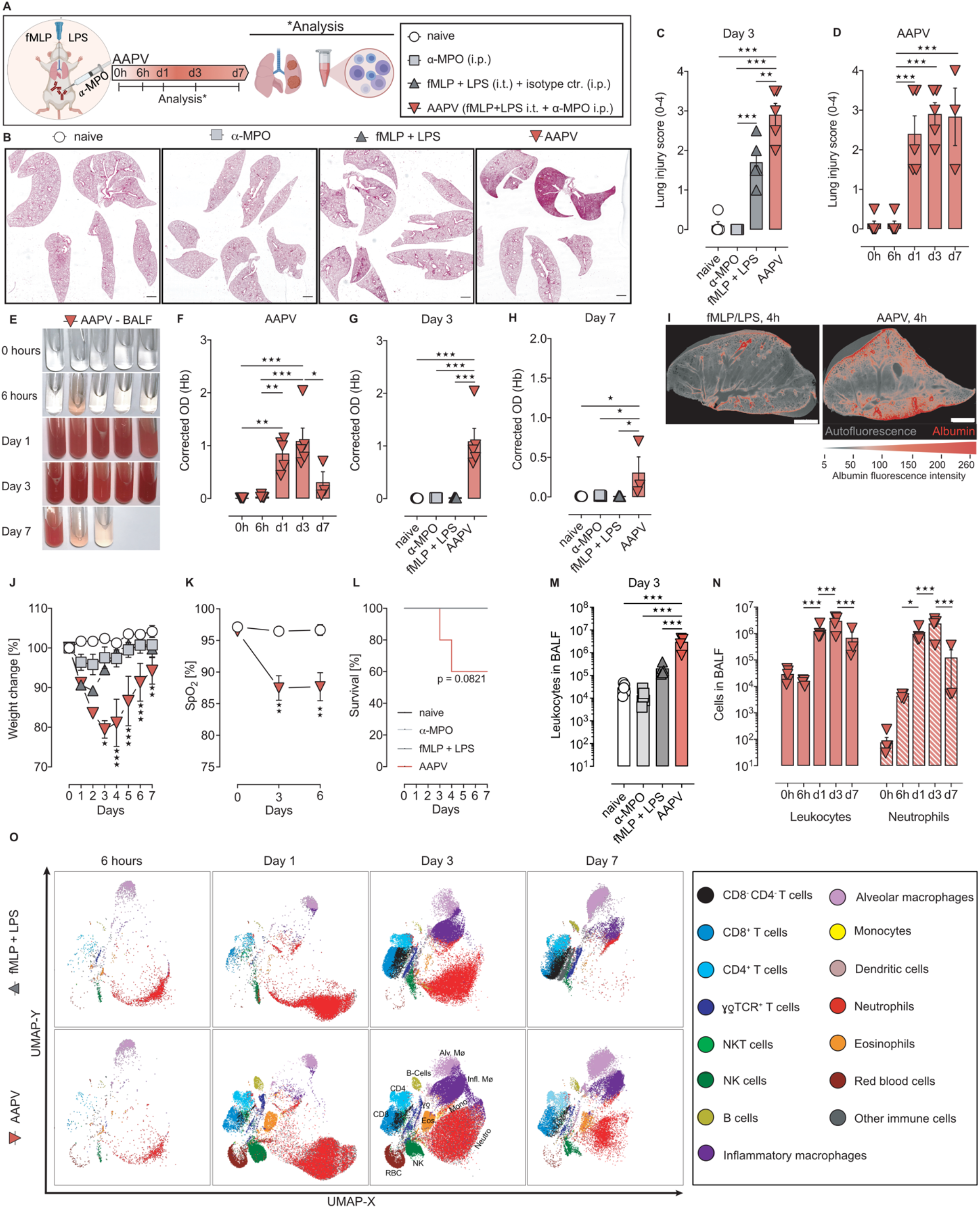
Anti-MPO antibodies and low dose bacterial ligands induce severe pulmonary vasculitis in a novel mouse model of pulmonary vasculitis. **(A)** Schematic of ANCA-associated pulmonary vasculitis (AAPV) induction. Anti MPO, monoclonal antibodies 6D1 and 6G4 **(B)** H&E stainings of lung cryosections 3d after disease induction from the indicated mouse groups. Bar = 500 µm. **(C, D)** Lung injury score (0-4) quantified from H&E lung cryosections. **(E-H)** Representative photographs (E) and quantification (F-H) of pulmonary hemorrhages in BALF. Treatments and time points as indicated. Hb, hemoglobin. **(I)** Light-sheet fluorescence microscope images from clarified left lungs isolated from mice treated as indicated and injected with Texas red-labeled albumin. Shown is a representative of two mice. **(J-L)** Kinetics of weight loss (J), SpO_2_(%) in peripheral blood (K), and survival (L) of mice as indicated in (A). **(M-O)** Flow cytometric quantification (M, N) and UMAP dimensionality reduction plots (O) of immune cells in BALF. Cell identity gates are shown in Fig. S2. Each symbol represents an individual mouse. Shown is mean ± S.E.M. Shown is a representative of at least three independent experiments with n=5 mice/group unless otherwise indicated. *, P<0.05; **, P<0.01; ***, P<0.001. AAPV, ANCA-associated pulmonary vasculitis; BALF, bronchoalveolar lavage fluid; OD, OD_400_ – OD_600_.

Systemic application of two murine anti-MPO monoclonal antibodies (Fayçal et al. 2022)(Fig. S1 A) into C57BL6/J mice failed to induce lung pathology (Fig. 2 B), consistent with two previously described with other polyclonal anti-MPO IgGs (Shochet, Holdsworth, and Kitching 2020) as well as in kidney, liver or skin (data not shown). Given that preceding bacterial often trigger infections ANCA-vasculitis episodes in patients (Timmeren, Heeringa, and Kallenberg 2014) by mobilizing and activating neutrophils (Huugen et al. 2005), we applied a combination of anti-MPO IgG together with low doses of bacterial ligands fMLP and LPS. Indeed, this caused marked pulmonary pathology characterized by large inflammatory cellular infiltrates and hemorrhages (Fig. 2 B and Fig. S1B and C). Severe and long-lasting lung damage was specific to the combined application, whereas single components did not suffice (Fig. 2 C and D, and Fig. S1 D). fMLP and LPS locally applied into the lung synergistically enhanced neutrophil mobilization (Fig. S1 E) and activation (Fig. S1 F), providing a mechanistic basis for the synergy with anti-MPO IgG for AAPV induction.

Concomitant with lung pathology, combination of bacterial ligands plus the monoclonal anti-MPO IgG antibodies resulted in hemorrhages in the lung (Fig. 2 E), mimicking the clinical hallmark of severe pulmonary vasculitis in patients (Jennette and Falk 1997). Blood accumulation in the bronchoalveolar space started as soon as 6 h after disease induction, peaked by day 3 and was partially resolved by day 7 (Fig. 2 E and F, and Fig. S1 G). Anti-MPO antibodies or bacterial ligands alone did not result in pulmonary hemorrhages (Fig. 2 G and H, and Fig. S1 H and I). Consistent with this, light-sheet microscopy of whole lungs showed widespread vessel leakage in the lungs of mice with active AAPV (Fig. 2 I). These findings were reflected by decreased mouse well-being (Fig. 2 J) and impaired lung function (Fig. 2 K), resulting in mortality of up to 40% of the mice (Fig. 2 L).

Combination of microbial ligands and anti-MPO monoclonal antibodies induced also a much stronger infiltration of immune cells into the bronchoalveolar space than the individual components on their own (Fig. 2 M and Fig. S1 J and K). Immune cell infiltration peaked on day 3 after vasculitis induction, with neutrophils comprising most of the cellular infiltrate and was strongly decreased by day 7 (Fig. 2 N and O), when pulmonary hemorrhages were resolving (Fig. 2 F).

In conclusion, we present here a novel and reproducible model of ANCA-associated pulmonary vasculitis that mimics clinical hallmarks of severe lung manifestations in patients.

### STING activation promotes autoimmune pulmonary vasculitis

Having identified activated cGAS in patients undergoing active vasculitis, we next investigated whether DNA sensing by the cGAS/STING pathway was important for pathogenesis in our mouse AAPV model. We observed extracellular DNA accumulation in the bronchoalveolar space of mice undergoing AAPV (Fig. 3 A) in a kinetics that closely correlated with the amount of blood in BAL (Fig. 2 F). This DNA had mostly a genomic rather than mitochondrial origin during the early phases of disease induction (Fig. 3 B). While fMLP/LPS treatment alone caused a similar increase in extracellular DNA concentration compared to mice undergoing AAPV (Fig. 3 A), it was clearly not sufficient to induce lung pathology (Fig. 2 B and G), highlighting the necessity of anti-MPO IgG for severe disease progression. Indeed, the anti-MPO antibodies used in this study were able to bind to the extracellular DNA released from *in vitro* activated neutrophils (Fig. 3 C), suggesting that DNA/MPO/IgG immune complexes played a pathogenic role.

**Figure 3.**
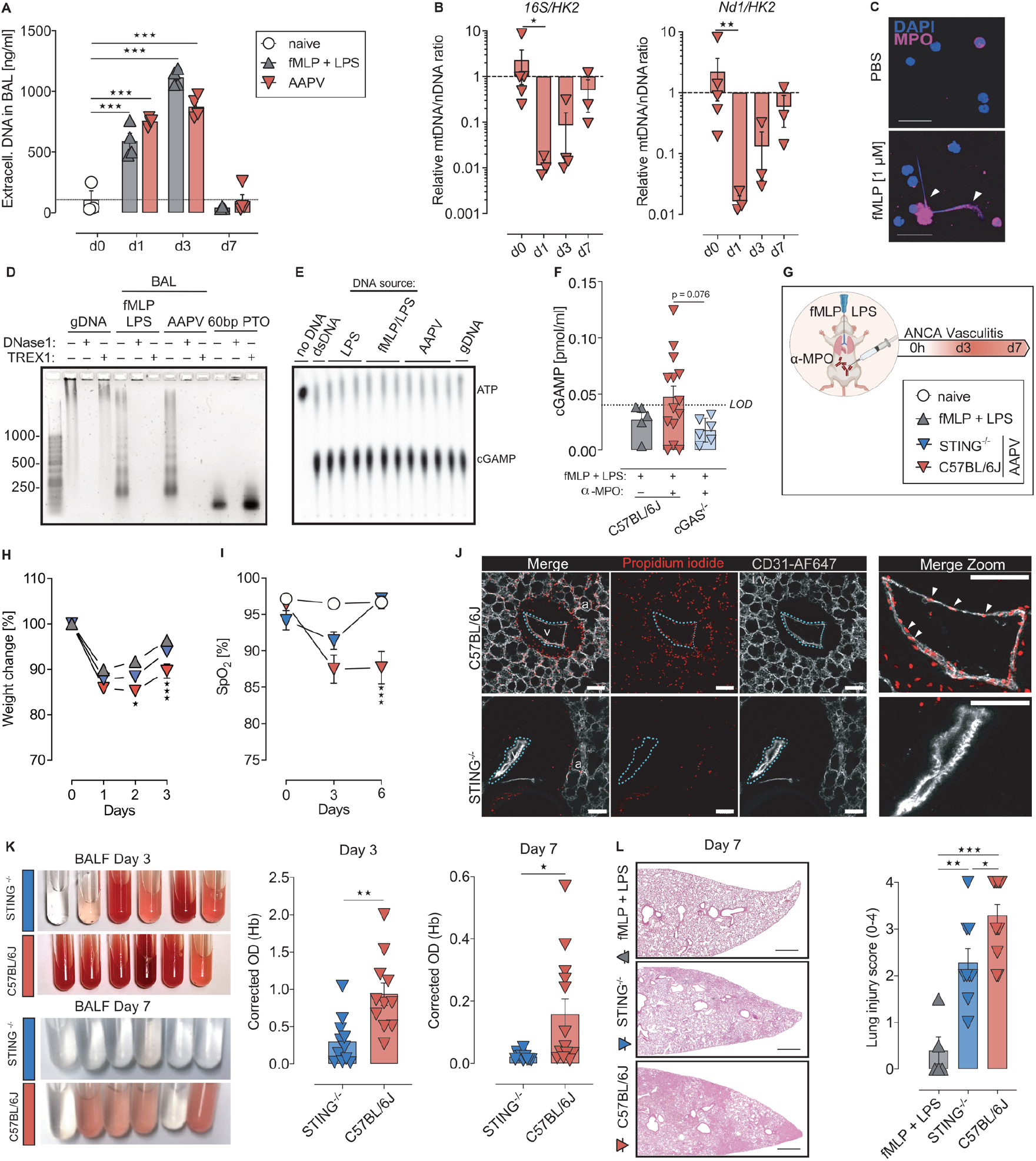
STING promotes progression of AAPV. **(A)** Cell-free DNA in the BALF of mice treated as indicated. Each dot represents an individual mouse. Data is representative of 2 experiments and 3-4 mice/group. **(B)** Cell-free mitochondrial (mt) DNA (*16S* or *Nd1*) to nuclear (n) DNA (*HK2*9) ratio obtained by qRT-PCR in BALF of AAPV mice at the indicated time points after disease induction. **(C)** Confocal microscopy images of neutrophils treated as indicated and stained with anti-MPO mAbs. Scale bar, 50 µm. Arrow heads indicate extracellular MPO-decorated DNA. **(D)** Gel electrophoresis of genomic DNA (gDNA), DNA isolated from BALF of mice 3 days after treatment as indicated, or 60mer TREX1-protected phosphorylated DNA treated or not with DNases *ex vivo* as indicated. Data is representative of 2 experiments with pooled DNA isolated from 3 mice per group. **(E)** cGAS activation assay using recombinant cGAS, ATP, and either dsDNA (lane 2), gDNA (lane 13), or DNA isolated from the BALF of mice treated 3 days earlier with LPS (lanes 3-5), fMLP/LPS (lanes 6-8) or fMLP/LPS + anti-MPO (lanes 9-12). Each lane represents an independent DNA sample. **(F)** cGAMP quantification in the BALF of mice treated as indicated 3 days earlier. Each dot represents a single mouse. LOD, limit of detection. **(G)** Schematic of experimental design in wild-type or Tmem173^gt/gt^ mice (STING^−/−^). **(H, I)** Kinetics of weight loss (H) and SpO_2_ [%] in peripheral blood (I) of mice treated as in (G). Shown is a representative from 3 (H) or 2 (I) experiments (n= 5 mice/group). **(J)** Confocal microscopy images of unfixed vibratome lung slices showing alveoli (a) and a medium-sized pulmonary blood vessel (v) stained *in vivo* with anti-CD31 and *ex vivo* with propidium iodide (PI). Bar = 100 µm. **(K)** Representative photographs and quantification of pulmonary hemorrhages in BALF. Treatments and time points as in (G). Shown is pooled data from 3 experiments. Hb, hemoglobin. **(L)** H&E staining of lung sections and lung injury score (0-4) for mice treated as indicated. Each dot is an individual mouse from 2 pooled experiments. Scale bar, 500 µm. Bars and line graphs represent mean ± SEM. *, P < 0.05; **, P < 0.01.***, P < 0.001.

Extracellular DNA fragments displaying characteristics of apoptotic DNA laddering were detected in the bronchoalveolar space of fMLP/LPS-treated or AAPV mice (Fig. 3 D), indicating release and partial degradation of DNA during inflammation. Despite its susceptibility to TREX1- and DNase1-mediated degradation (Fig. 3 D), the released DNA had the capacity to activate the DNA sensor cGAS as demonstrated by cGAMP generation using a sensitive cell-free assay with recombinant cGAS and ^32^P-labelled ATP (Civril et al. 2013), independently of anti-MPO administration (Fig. 3 E). These data indicate that extracellular immunogenic DNA accumulated in the lung of mice treated with bacterial ligands and that anti-MPO antibodies were required for disease progression, suggesting that DNA/MPO/IgG complexes were required to deliver extracellular immunogenic DNA to the cytosol for cGAS activation.

Consistent with our findings in patients (Fig. 1A), we observed that 50% of mice in the AAPV group had cGAMP levels above detection level, whereas none of the mice treated only with bacterial ligands did (Fig. 3 F), supporting *in vivo* cGAS activation during AAPV. To validate the contribution of cGAS/STING pathway, we compared vasculitis induction in wild-type (C57BL/6J) and Tmem173^gt/gt^ mice expressing non-functional STING (here referred to as STING^−/−^ mice) (Fig. 3 G). Deficiency in functional STING prevented the AAPV-induced weight loss observed in WT mice (Fig. 3 H) and restored lung function 6 days after disease induction compared to WT mice, which still showed decreased blood oxygenation at that time point (Fig. 3 I). Consistent with these findings, functional STING promoted endothelial cell death during AAPV induction (Fig. 3 J), leading to more severe pulmonary hemorrhages (Fig. 3 K) and lung histopathology (Fig. 3 L). We thus conclude that DNA recognition via the cGAS/STING pathway promoted lung injury and hemorrhages resulting in decreased pulmonary function.

### Type-I IFN is required for ANCA pulmonary vasculitis

STING activation following cGAS-mediated DNA recognition generally results in signaling via TBK1/IRF3 and IFN-β expression (Ishikawa and Barber 2008; Motwani, Pesiridis, and Fitzgerald 2019). Consistent with our findings of increased IFN-I signature in patients (Fig. 1B-D), we hypothesized that type-I IFN (IFN-I) plays a pathological role in ANCA vasculitis. Indeed, we observed increased *Ifnβ* expression (Fig. 4 A), and IFN-I signature at the peak of vasculitis as demonstrated by the upregulation of different interferon stimulated genes (ISGs) in the lung of mice with AAPV (Fig. 4 B and S3 A). STING^−/−^ mice expressed lower pulmonary *Ifnβ* levels after vasculitis induction than their wild-type counterparts (Fig. 4 C), indicating that STING promoted *Ifnβ* production. Consequently, STING^−/−^ mice showed reduced ISGs expression after induction of pulmonary vasculitis (Fig. 4 D). Within the analyzed ISGs, we found also *Oasl1, Cxcl10*, and *Mx-1* to be increased during disease progression (Fig. S3 A), albeit in a STING-independent manner (Fig. S3 B), likely due to IFN-I-unrelated inflammatory cues (Schneider, Chevillotte, and Rice 2014).

**Figure 4.**
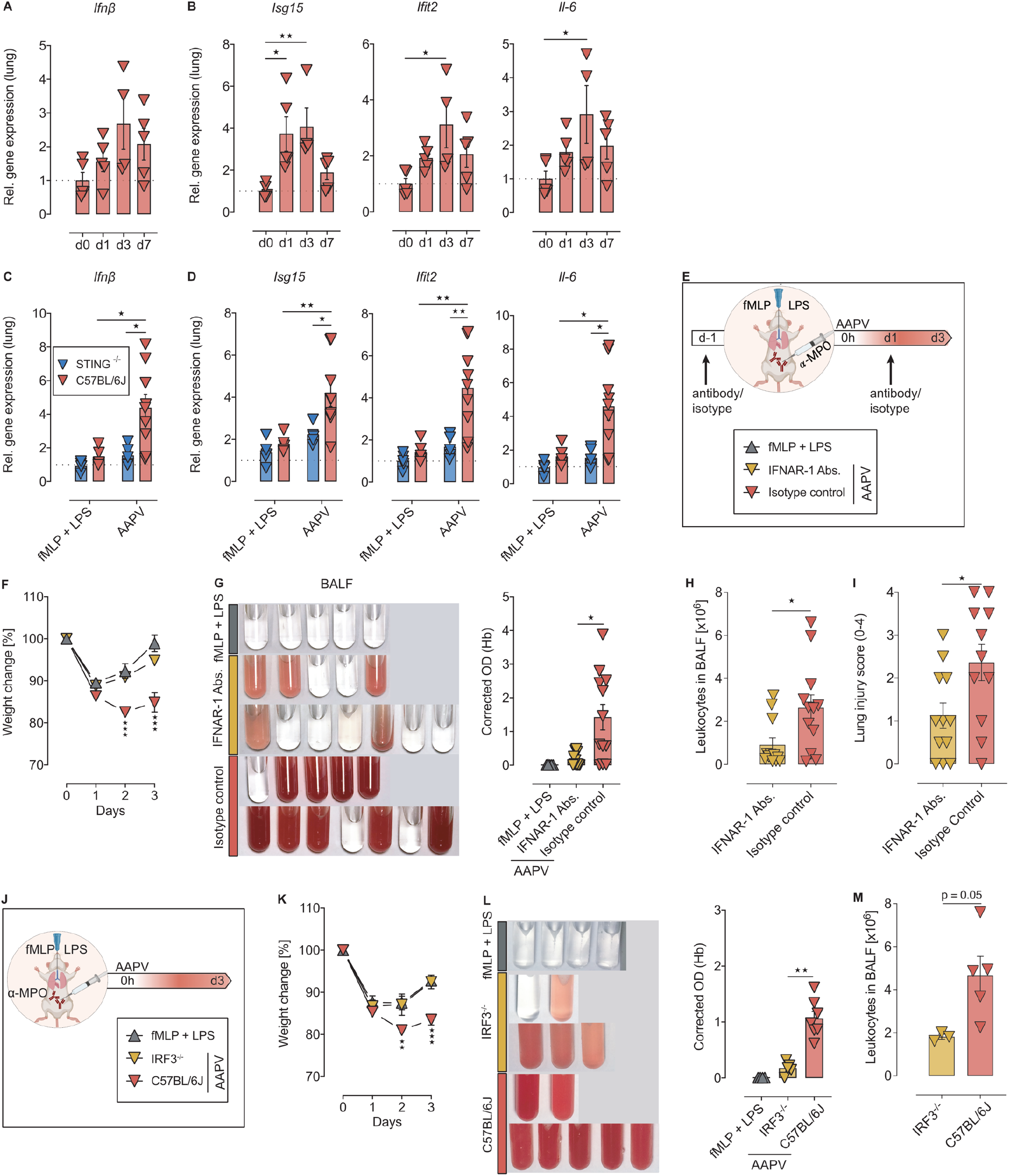
IFNAR-1 and IRF3 promote severe anti-MPO-associated pulmonary vasculitis. **(A-D)** Relative expression of *Ifnβ* gene (A, C) and indicated ISGs (B, D) at different time points after AAPV induction in WT mice (A, B) or on day 3 in the indicated mice (C, D). Each dot is an individual mouse. Data are representative of 2 experiments. **(E)** Schematic representation of experiment with IFNAR-1 blocking antibodies. **(F-I)** Weight loss (F), hemorrhages in BALF (G), leukocyte counts (H) and lung injury score (I) in WT mice treated as indicated. Each dot is an individual mouse. Shown are 2 pooled experiments. **(J)** Schematic representation of experiments with IRF3^−/−^ mice. **(K-M)** Weight loss (K), hemorrhages in BALF (L), and leukocyte counts (M) in WT and IRF3^-/-^ mice treated as indicated. Each dot is an individual mouse. Shown are results from 2 pooled experiments. Bars and line graphs represent mean ± SEM. *, P < 0.05; **, P < 0.01; ***, P < 0.001. Hb, hemoglobin.

Antibody-mediated blockage of IFNAR-1 (Fig. 4 E) diminished AAPV progression as demonstrated by reduced weight loss (Fig. 4 F), lower blood content in the BAL (Fig. 4 G) and leukocyte infiltration (Fig. 4 H) as well as decreased lung pathology (Fig. 4 I) compared to isotype-treated mice at the peak of disease progression (day 3), indicating a crucial role for the STING/IFN-I axis on progression towards severe AAPV. To obtain further independent evidence for the role of STING/IFN-I axis, we employed mice lacking the transcription factor IRF3 (Fig. 4 J), which is required to transduce STING signaling into IFN-I production (Ishikawa and Barber, 2008). As in wild-type mice treated with blocking α-IFNAR-1 antibodies, IRF3^−/−^ mice showed less weight loss (Fig. 4 K), pulmonary hemorrhages (Fig. 4 L) and leukocyte infiltration (Fig. 4 M) upon challenge with bacterial ligands and anti-MPO IgG.

Thus, we conclude that STING activation promotes an IRF3-dependent IFN-I signature that is required for the development of severe AAPV.

### Functional dichotomy between resident and inflammatory macrophages in ANCA vasculitis progression

Having identified STING/IFN-I as a major driver for severe AAPV, we next investigated the cellular source of IFN-β, by using reporter mice in which transcriptional activation of *Ifnβ* results in luciferase production (Fig. 5 A). Consistent with our findings identifying an IFN-I signature in patients and mice (Fig. 1 B and C, and Fig. 4 A and B), administration of bacterial ligands and anti-MPO IgG induced luciferase expression in the lung, further demonstrating *Ifnβ* transcription during AAPV (Fig. 5 B). Bioluminescent *ex vivo* analysis of sorted cells indicated that most IFN-β was derived from inflammatory macrophages, whereas lung-resident alveolar macrophages, lymphoid cells, neutrophils, eosinophils, epithelial cells and endothelial cells produced little or no detectable IFN-β (Fig. 5 C).

**Figure 5.**
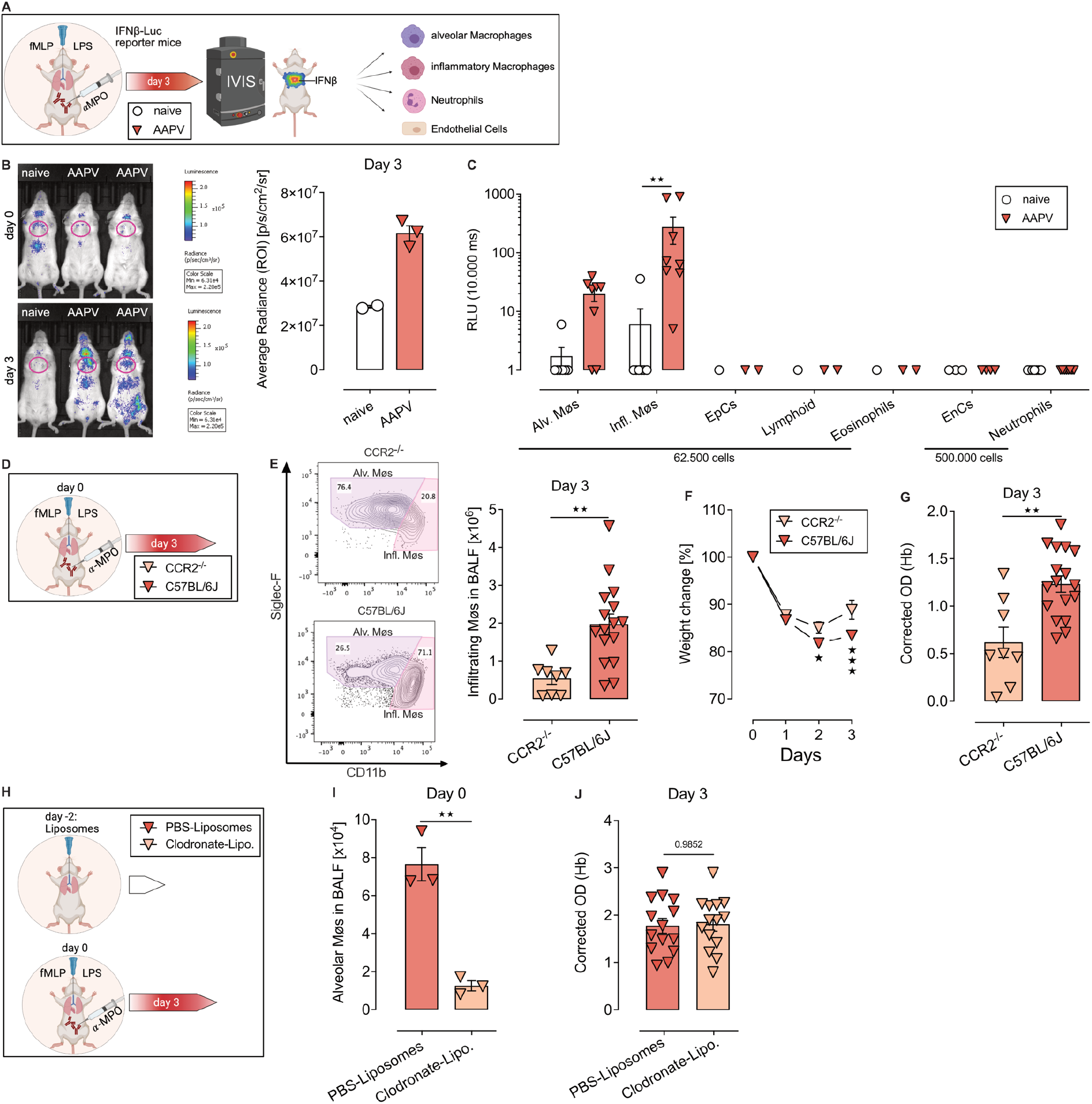
Infiltrating macrophages produce IFN-I and enhance severity of AAPV. **(A)** Schematic representation to measure *Ifnβ* promoter activity in reporter mice by *in vivo* and *ex vivo* bioluminescence imaging. **(B)** Representative photographs of *Ifnβ*-Luc reporter mice treated as indicated, and bioluminescence quantification of the region of interest (ROI, red gate). Shown in one of three experiments. **(C)** ex vivo bioluminescence quantification of FACS-sorted cell populations (gates in Fig. S2 and S4A). Each dot represents cells from an individual mouse. Shown is pooled data from 3 (Møs, Encs and neutrophils) or 2 (all other cell types) experiments. Alv, alveolar; infl, infiltrating; Møs, macrophages; EpCs, epithelial cells; EnCs, endothelial cells. **(D)** Schematic representation of AAPV induction in mice. **(E)** Flow cytometric plot showing percentages of Alv. and Infl. Møs gated as in Fig. S2., and respective quantification. Shown is a representative of two independent experiments with n=8-17 mice/group **(F, G)** Weight loss (F) and hemorrhages in BALF (G) in indicated mice undergoing AAPV. **(H)** Schematics of treatment with liposomes before onset of AAPV. Lipo, liposomes. **(I)** Quantification of Alv. Møs on the day of AAPV induction. Each dot represents an individual mouse from a representative of two independent experiments with n=3 mice/group **(J)** Quantification of hemorrhages in BALF at the peak of AAPV severity (day 3). Each dot represents an individual mouse from a representative of two independent experiments with n=7 mice/group. Bar and line graphs show mean ± SEM. *, P < 0.05; **, P < 0.01; ***, P < 0.001. Hb, hemoglobin.

To elucidate the role of *Ifnβ*-producing inflammatory macrophages on AAPV progression, we administered bacterial ligands as well as anti-MPO IgG in mice lacking *Ccr2* or not (Fig. 5 D). Inflammatory infiltrating macrophages were of monocytic origin as indicated by Ly-6C and F4/80 expression (Fig. S4 B) as well as dependency on Ccr2 expression (Fig. 5 E). Ccr2-dependent inflammatory macrophages promoted disease severity as indicated by less pronounced weight loss (Fig. 5 F) and pulmonary hemorrhages (Fig. 5 G) in *Ccr2^−/−^* mice compared to their Ccr2-competent counterparts.

Lung-resident alveolar macrophages (AM) also produced IFN-β, albeit to a much lower level than infiltrating macrophages (Fig. 5 C). To investigate the contribution of AMs, we used clodronate liposomes to deplete them 2 days before disease induction (Fig. 5 H). Clodronate liposomes significantly reduced AM numbers at the time of disease induction (Fig. 5 I) without affecting the extent of pulmonary hemorrhages (Fig. 5 J) at the peak of disease (day 3), suggesting they did not play a pathogenic role in the initial phase of disease progression.

However, when we analyzed BALF infiltrates by cytology, we found large mononuclear cells having internalized RBCs (Fig. 6 A). Interestingly, these hemophagocytes were absent in the BALF of AM-depleted mice (Fig. 6 A). Iron staining revealed that indeed AMs, but not inflammatory macrophages or neutrophils, took up extravascular RBCs (Fig. S4 C).

**Figure 6.**
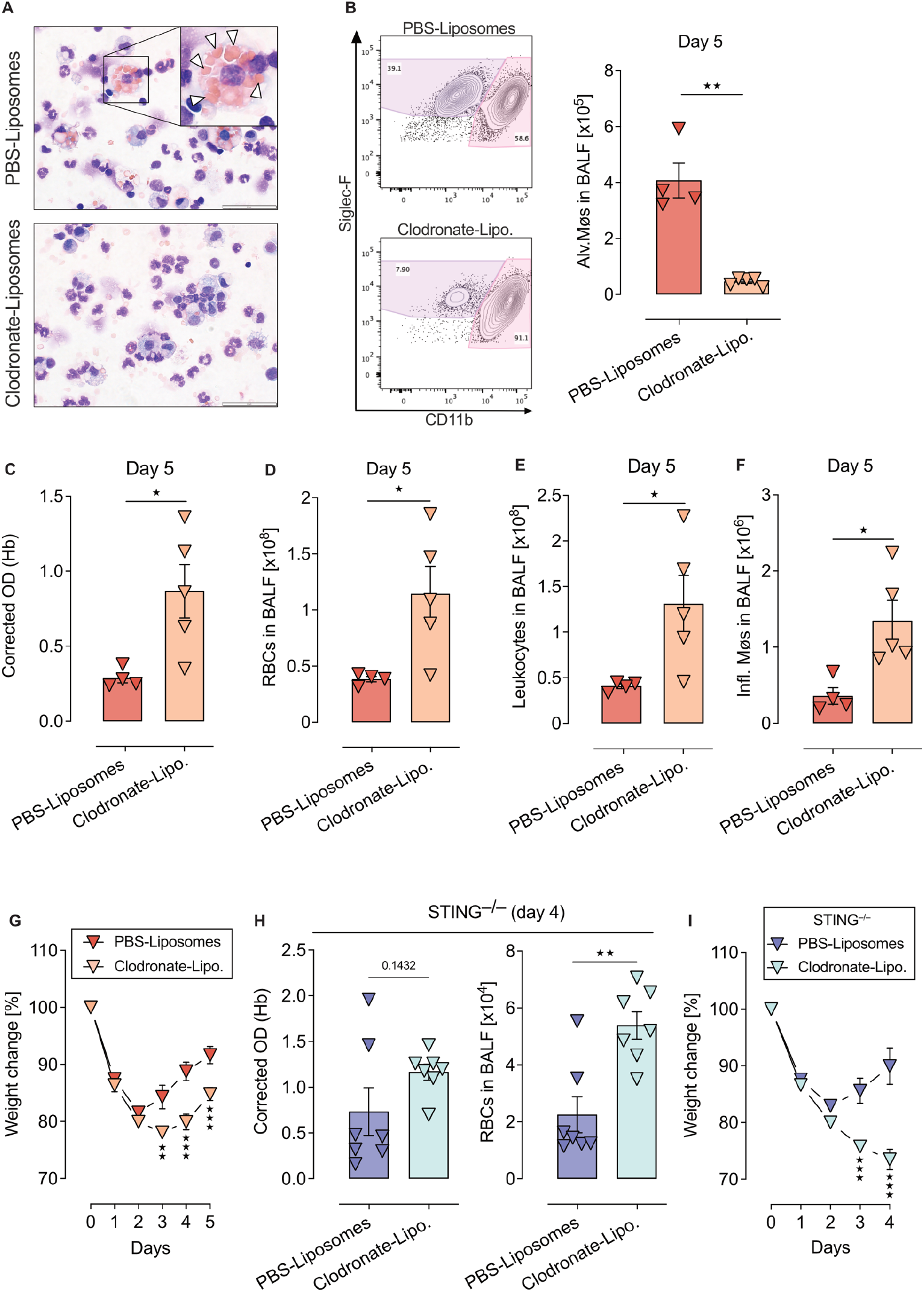
Alveolar macrophages promote homeostasis by clearing extravascular RBC. **(A)** Giemsa staining of BALF cytospins from mice treated with PBS- or clodronate liposomes. White arrow heads indicate engulfed RBCs. **(B-G)** Quantification of Alv. Møs (B), hemorrhages in BALF (C, D), Leukocytes (E), infiltrating Møs (F), and weight loss (G) in BL6 mice during the recovery phase (5 days after APPV induction) and treated with liposomes before disease induction as in Fig 5H. Each symbol represents an individual mouse. Shown is a representative of three experiments with n=7 mice/group **(H, I)** Hemorrhages in BALF (H) and weight loss (I) in STING^−/−^ mice undergoing AAPV and treated with liposomes as in (B-G). Each symbol represents an individual mouse. Shown is pooled data from two independent experiments. Bar and line graphs show mean ± SEM.*, P < 0.05; **, P < 0.01; ***, P < 0.001. Hb, hemoglobin.

We therefore hypothesized that AM promoted lung homeostasis rather than disease progression by clearing extravascular RBCs arising from pulmonary hemorrhages and, thereby, reducing extracellular heme, a well-known proinflammatory mediator (Consonni et al. 2021; Weis et al. 2017). To investigate this, we analyzed disease recovery in AM-depleted mice (as in Fig. 5 H) five days after disease onset, when hemorrhages were being cleared (Fig. 2 E and F). AM were still strongly depleted in clodronate-liposome-treated mice at that time point (Fig. 6 B). While control mice had cleared most of the pulmonary hemorrhages by day 5, AM-deficient mice showed significantly higher hemorrhages in BALF (Fig. 6 C and D). Consistent with extravascular heme being a proinflammatory stimulus, AM-depleted mice that were compromised in RBC clearance showed significantly higher leukocyte infiltration into the lung (Fig. 6 E), including inflammatory macrophages (Fig. 6 F), and weight loss (Fig. 6 G) during the recovery phase.

To further confirm that clearance of RBCs is necessary for reverting to homeostasis following AAPV induction, we depleted AMs in STING-deficient mice, which showed a faster recovery than wild-type counterparts (Fig. 3 H and I, K and L). In line with our previous findings, AM depletion in STING^−/−^ mice also resulted in increased hemoglobin levels and RBCs in BALF (Fig. 6 H). Interestingly, the protective effect of STING-deficiency was abrogated by AM depletion (Fig. 6 I), indicating that extravascular RBC accumulation prevented recovery from disease even in the absence of functional STING.

Altogether these data indicate a functional dichotomy of lung macrophages subpopulations in AAPV. While inflammatory macrophages infiltrate the lung in a Ccr2-dependent fashion and produce pathogenic IFN-β, lung-resident alveolar macrophages counteracted disease progression by clearing extravascular RBCs and reducing the number of inflammatory macrophages.

### Pharmacological inhibition of the STING/IFNAR pathway ameliorates autoimmune pulmonary vasculitis

Based on our results, we next explored the therapeutic efficacy of pharmacologically targeting STING or IFNAR-1 signal transduction with established small-molecule inhibitors. STING inactivation by the covalent small-molecule inhibitor H151 (Fig. 7 A) prevented vasculitis-associated weight loss (Fig. 7 B) and reduced the incidence of mice with pulmonary hemorrhages (Fig. 7 C). Furthermore, H151-treated mice showed lower hemorrhages (Fig. 7 D) and leukocyte infiltration (Fig. 7 E) in the BAL, although the lung injury score, which accounts for non-hemorrhagic lung alterations, was not significantly ameliorated (Fig. 7 F).

**Figure 7.**
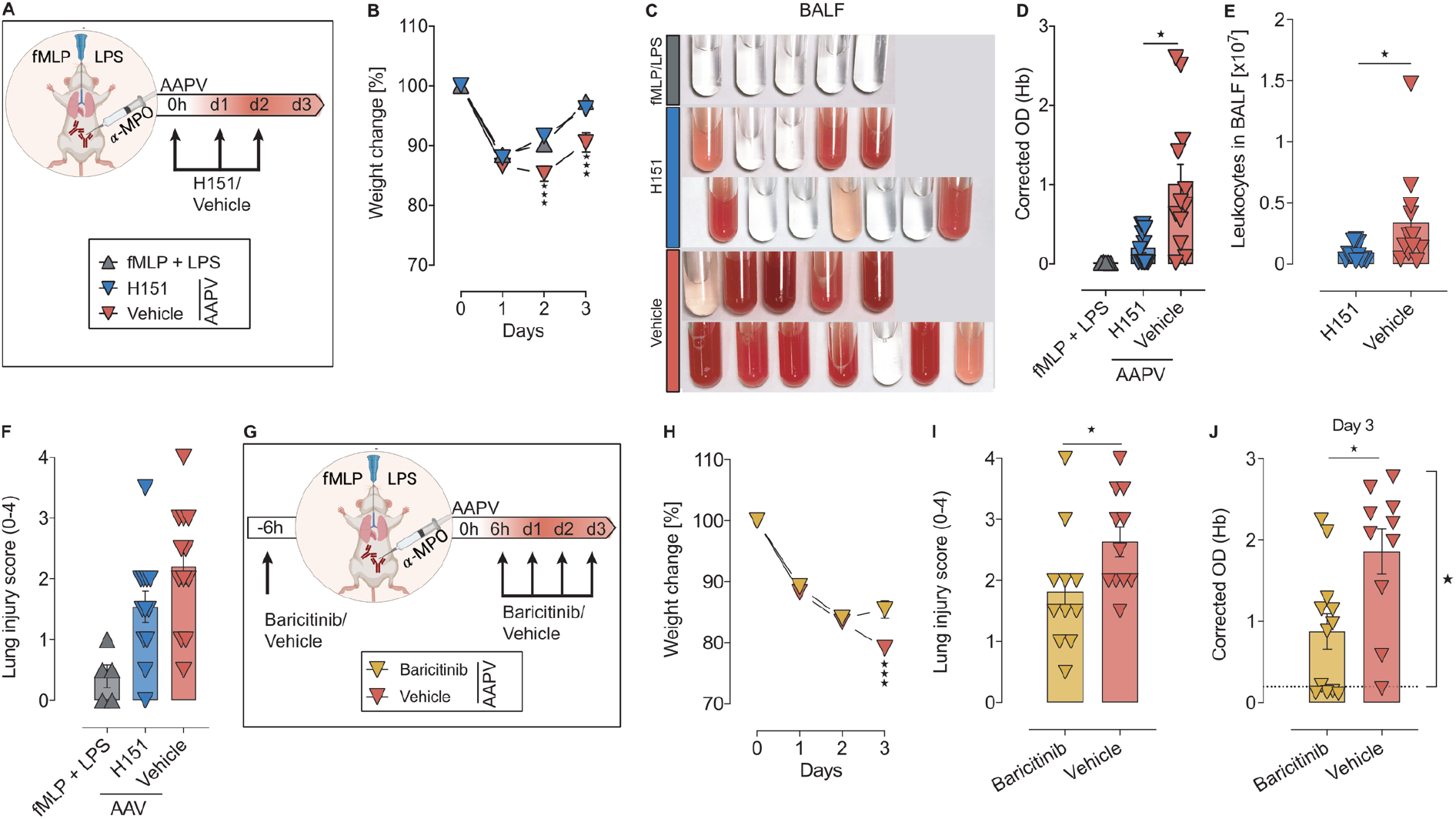
Pharmacological inhibition of STING or JAK/STAT pathways ameliorates AAPV in mice. **(A)** Schematic representation of experiment with H151 STING inhibitor. **(B-D)** Weight loss kinetics (B), hemorrhages in BALF (C, D), leukocyte infiltration (E) and lung injury score (F) in mice treated as in (A). **(G)** Schematic representation of experiment with JAK1/2 inhibitor baricitinib. **(H-J)** Weight loss (H), lung injury score (I) and hemorrhages in the BALF (J) of WT mice treated as in (A). Each dot is an individual mouse (D-F, I, J). Bar and line graphs show mean ± SEM. Data is pooled from 2 independent experiments. *, P < 0.05; **, P < 0.01.***, P < 0.001. Hb, hemoglobin.

Since IFNAR-1 was required for the induction of severe pulmonary vasculitis, we hypothesized that the small-molecule inhibitor baricitinib, which targets JAK1 downstream of IFNAR-I and is used to treat interferonopathies in patients (Sanchez et al. 2018), may ameliorate AAPV (Fig. 7 G). Baricitinib treatment reduced weight loss (Fig. 7 H) and lung histopathology (Fig. 7 I) compared to vehicle treatment. Importantly, 10 of 11 (90.9%) vehicle-treated mice developed blood in the BAL, whereas baricitinib treatment resulted in 8 of 13 (61.5%) mice developing hemorrhages at day 3 after disease induction, and to a lower degree (Fig. 7 J).

In conclusion, pharmacological inhibition of the STING and JAK1/2 pathways with well-characterized small-molecule inhibitors improved general well-being and lung hemorrhages in our mouse model for AAPV.

## Discussion

AAV is a group of autoimmune diseases affecting small- and medium-sized blood vessels with a mortality rate of up to 80% within the first year of diagnosis if left untreated (Córdova-Sánchez et al. 2016). Current therapies rely on systemic steroid administration with severe side effects, accounting for about half of the mortality observed during the first year of treatment, mostly due to infections (Little et al. 2010). Thus, there is an urgent need to identify specific effector mechanisms that can be therapeutically targeted. Here, we report that patients with active autoimmune AAV showed increased cGAMP levels and IFN-I signature, suggesting pathogenic DNA recognition by the cGAS/STING axis. Using a novel mouse model for ANCA pulmonary autoimmune vasculitis, we demonstrate that the STING/IRF3/IFN-I axis is required for severe pulmonary bleeding, immune cell infiltration and lung dysfunction, which are all hallmarks of the most life-threatening form of AAV in patients (Jennette and Falk 1997). Pharmacological inhibition of STING, IFNAR-1 or JAK1/2 decreased disease severity and accelerated recovery. Thus, our results identify STING and IFN-I as important mediators of disease progression, and open potential therapeutic venues by targeting those pathways.

The mouse model here presented offers several distinctive advantages over currently available models. It is easily induced and based on defined mouse monoclonal antibodies directed against murine MPO rather than ill-defined polyclonal antibodies or protein-based immunization, which may cause highly variable antibody titers within the experiment groups (Shochet, Holdsworth, and Kitching 2020). In addition, this experimental model is unique in inducing pulmonary disease, often occurring in life-threatening AAV. Progression of pulmonary vasculitis depended on the combined application of anti-MPO monoclonal antibodies together with low doses of the bacterial ligands fMLP and LPS to model bacterial infections often observed preceding AAV flares (Timmeren, Heeringa, and Kallenberg 2014). Low doses of fMLP and LPS synergized to enhance neutrophil recruitment and activation resulting in severe AAPV upon anti-MPO administration, consistent with the role of these cells during disease progression (Xiao et al. 2005).

STING activation results in pathogenic IFN-I levels in a Trex-1 deficiency heart injury model (Haag et al. 2018) and in a murine model for COVID-19 (Domizio et al. 2022). We demonstrate that a covalent small-molecule inhibitor of human and mouse STING can reduce disease severity and pulmonary hemorrhages in the ANCA vasculitis mouse model. STING is activated following cGAS-mediated DNA recognition in the cytosol (Ablasser, Goldeck, et al. 2013; Diner et al. 2013; Sun et al. 2013; Wu et al. 2013). We provide evidence in mice and humans that DNA recognition by cGAS/STING/IFN-I axis promotes AAV progression. Albeit detected at low levels, cGAMP concentration was increased in the BAL of diseased mice and PBMCs of patients with active AAV. Low cGAMP concentrations may be the consequence of its ability to spread intercellularly, thereby amplifying the reach of the inflammatory response (Ablasser, Schmid-Burgk, et al. 2013; Zhou et al. 2020). Nevertheless, these low levels were sufficient to promote STING activation and disease progression.

Although extracellular DNA is known to reach the cytosol and activate cGAS/STING *in vivo* (Gehrke et al. 2013), the mechanisms driving its translocation into the cytosol remained unclear. Delivery of extracellular DNA into the cytosol has also been recently proposed in a non-autoimmune model of STING-induced aortic aneurism and dissection (Luo et al. 2020). High-dose LPS alone has recently been shown to induce rapid mtDNA release resulting in cGAS/STING-mediated endothelial cell damage in mouse lungs (Huang et al. 2020). However, these findings cannot explain the vasculitis progression reported in our study using low dose LPS because disease depended on anti-MPO administration. fMLP/LPS was sufficient to mediate the release of cGAS-stimulating DNA. However, pulmonary hemorrhages depended on anti-MPO administration, which is consistent with increased circulating levels of DNA/MPO complexes in patients with active AAV (Söderberg et al. 2015). Therefore, we hypothesize that immune complexes formed between MPO-decorated DNA released by activated neutrophils and anti-MPO IgGs promote the internalization of immunogenic DNA. In support of this, Fc receptors contribute to the cytosolic translocation of DNA/α-dsDNA immune complexes (Shin et al. 2013). Further experiments are required to unveil how exactly extracellular DNA is shuttled into the cytosol for cGAS/STING recognition.

STING signals via the TBK1/IRF3 or the canonical NF-*κ*B pathways to induce expression of pro-inflammatory mediators such as IFN-I and TNFα, respectively (Motwani, Pesiridis, and Fitzgerald 2019). IFN-I is at the center stage of congenic (Kretschmer and Lee-Kirsch 2017), but also spontaneous autoimmune diseases such as SLE (Banchereau and Pascual 2006). We have shown in two independent approaches that the TBK-1/IRF3/IFN-I pathway promotes AAPV development. In IRF3^−/−^ mice and in wild-type mice treated with blocking antibodies directed against IFNAR-1, the development of vasculitis in the lung was significantly attenuated. This was in clear contrast with a mouse model for SAVI caused by a self-activating mutation in STING, where IRF3- or IFNAR1-deficiency did not protect from disease development (Luksch et al. 2019; Warner et al. 2017), suggesting that different pathophysiological mechanisms drive acquired AAPV and congenic SAVI in the lung.

Our results show that both in patients and mice, ANCA vasculitis is associated with an IFN-I signature. The role of IFN-I in the pathogenesis of ANCA-vasculitis has been, however, controversial. ISG or IFN-I have been shown to be increased in patients with active vasculitis (Kessenbrock et al. 2009). However, a recent study did not find increased IFN-I or ISG expression in ANCA vasculitis patients (Batten et al. 2021). The reasons for this apparent disagreement are unresolved, although differences in disease severity across the different patient cohorts cannot be excluded. Our results not only show an IFN-I signature in patients with ANCA-associated vasculitis but demonstrate that the STING/IRF3/IFN-I is pathogenic and blocking it at different levels ameliorates disease.

Inflammation is characterized by dynamic changes in the composition or organ-resident macrophages and infiltrating macrophages (Hou et al. 2021; Calum C. Bain and MacDonald 2022). Whereas the ontogeny of such populations is well characterized (Geissmann et al. 2010; Guilliams et al. 2014; Hume, Irvine, and Pridans 2019; C C Bain et al. 2012), whether they differentially contribute to inflammation remains unresolved. In the lung, it has been postulated that resident AMs are anti-inflammatory mostly due to production of immunosuppressive mediators such as TFGβ (Garbi and Lambrecht 2017), whereas infiltrating macrophages promote inflammation by release of cytokines such as TNF-*α* (Lin et al. 2008, 2) or TRAIL (Herold et al. 2008). Here, we demonstrate a clear functional dichotomy in lung macrophage subsets. Most IFN-β was produced by infiltrating macrophages and these cells promoted pulmonary hemorrhages during ANCA-mediated vasculitis, consistent with a pro-inflammatory function. On the other hand, tissue-resident alveolar macrophages did not modify AAPV progression in the early phases of disease induction, but accelerated recovery by clearing extracellular red blood cells and dampened inflammation, likely by reducing extracellular heme accumulation (Consonni et al. 2021). Of note, although extravascular RBC accumulated in large numbers in the lung of mice undergoing AAPV, we did not detect RBC intake by other phagocytes such as neutrophils, monocytes, and monocyte-derived infiltrating macrophages.

We are aware of some limitations in our study. We have focused on the clinically relevant effector phase of vasculitis and did not address the mechanisms leading to the formation of MPO-specific autoantibodies. Unveiling whether this depends also on STING activation was not within the scope of the present study. Future studies using other vasculitis models may clarify whether STING signaling also feeds back on the generation of pathogenic anti-MPO antibodies.

In summary, we established and used a novel mouse model for anti-MPO-associated vasculitis to show that genetic or pharmacological interference with the cGAS/STING/IFN-I axis significantly reduced disease severity. The elevated cGAMP and IFN-I signature observed in patients with active vasculitis suggests that targeting those pathways may be of therapeutic value.

## Materials and Methods

### Clinical study design

#### Cohort I

AAV patients were recruited from a single rheumatology tertiary center. Clinical details are described in Supplemental Table 1. Written informed consent was obtained from all subjects according to the Declaration of Helsinki and approval by the Institutional Review Board of the University of Bonn (210/19).

#### Cohort II

Patients were recruited from a specialist tertiary center during active disease with ANCA vasculitis. Enrolled patients were required to be on <10 mg prednisolone and on no other concurrent immunosuppression for a minimum of 3 months prior to enrollment. Active disease was assessed in patients with ANCA vasculitis using the Birmingham Vasculitis Activity Score (BVAS). Prior to treatment escalation, 100 ml blood samples were collected. Healthy controls were age and sex matched where possible. Ethical approval was obtained from the Cambridgeshire Regional Ethics Committee (REC08/H0306/21). All patients provided written informed consent. Patient details are described in Supplemental Table 2.

### Mice

C57BL/6J, *Tmem173*^gt/gt^ (referred here as STING^−/−^) (Sauer et al. 2011), *Mb21d1*^−/−^ (referred here as cGAS^−/−^) (Schoggins et al. 2014) *Irf3*^−/−^ (Sato et al. 2000) and Ccr2^-/-^ (Kurihara et al. 1997) mice were purchased from The Jackson Laboratory. IFN-β reporter mice (Lienenklaus et al. 2009) were kindly provided by Rayk Behrendt. All mice were bred on C57BL/6J background and maintained under SPF conditions at the animal facilities of the University Hospital Bonn. Female mice were used at 8-12 weeks of age. Mice were not randomized in cages, but each cage was randomly assigned to a treatment group. All animal experiments were approved by the corresponding government authority (Landesamt für Natur, Umwelt und Verbraucherschutz, NRW).

### Generation of anti-MPO monoclonal antibodies

Mouse monoclonal antibodies 6D1 (IgG2b) and 6G4 (IgG2c) directed against murine myeloperoxidase (MPO) were generated in exactly the same way as described for anti-MPO antibody clone 8F4 (van Leeuwen et al. 2008). Bulk production of 6D1 and 6G4 was performed by BioXcell.

### Pulmonary vasculitis model

Pulmonary vasculitis was induced by intratracheal (i.t.) application of 50 µl PBS containing 10 µg LPS (Serotype 026:B6, Sigma) and 10 µg fMLP (Sigma) with the help of a small-animal laryngoscope and mechanical ventilation (Harvard Apparatus) for 1 minute at 250 µl air/stroke and 250 strokes/minute as previously described (Holland et al. 2017), followed by intraperitoneal (i.p.) injection of 1 mg anti-MPO (clones 6D1 and 6G4, 0.5 mg of each clone) in 200 µl PBS. Control groups were either left untreated (naive) or received either anti-MPO (i.p.) alone or fMLP + LPS + isotype control (clones MPOC-137, MPC-11) in PBS (i.t.). Unless stated otherwise, analysis was performed on day 3 or 7 after induction. Weight loss was monitored daily.

### Pharmacological inhibition of IFNAR-1, STING and JAK/STAT signaling

IFNAR-1 was blocked as previously described by injecting mice with 250 µg α-IFNAR-1 (clone MAR1-5A3) antibody i.p. one day prior to vasculitis induction and then every second day. Control mice were treated equally but with an irrelevant isotype control (clone MOPC-21).

STING activation was inhibited as previously described (Haag et al. 2018) by injecting mice i.p. daily with 750 nM H151 (inh-h151, InvivoGen) starting on the day of AAPV induction. Control mice received vehicle only.

JAK/STAT pathway was inhibited as previously described (Carfagna et al. 2018) by oral gavage of 10 mg/kg body weight Baricitnib (Olumiant^®^, Lilly). Control mice received an equal amount of only vehicle by oral gavage.

### Depletion of alveolar macrophages *in vivo*

To deplete alveolar macrophages, 50 µl of Clodronate-or PBS-loaded liposomes (Control) were applied intratracheally two days prior to vasculitis induction under 2% isoflurane anesthesia.

### Pulse Oximetry

Blood SpO_2_ (%) was measured with the MouseOx^TM^ Pulse-oximeter (Starr Life Science). Mice were anesthetized with 2% isoflurane in air and the oximeter sensor was placed on the upper part of the shaved hind-leg. Once stabilized over a maximum of 3 minutes, SpO_2_ (%) was measured each second over 3-5 min and values were averaged. Data shown is the mean of 3 individual averaged measurements per animal.

### *In vivo* bioluminescence quantification

To quantify luminescence of *Ifnβ*-Luc reporter mice, we injected 4.5 mg of Luciferin per mouse i.p. Mice were then anesthetized with isoflurane and placed in the IVIS (Lumina LT Series III) machine 5 min later at 37°C. Luminescence was sequentially acquired ranging from 5 minutes to 1 sec of exposure time and a time point equal to all mice on the linear dynamic range was selected for quantification.

### Blood cell collection, BALF collection and lung fixation

Mice were killed by an overdose of Rompun/Ketamine. Mouse blood was collected from the lower aorta into heparinized capillaries. Red blood cells were lysed by incubating for 7 min at room temperature in PBS containing 10 mM NH_4_Cl, 0.1 mM EDTA and 0.01 M NaHCO_3_ (pH 7.3). Cells were centrifuged at 1500 rpm for 10 min at 4°C and the cell pellet resuspended in FACS buffer (PBS with 3% heat-inactivated fetal calf serum).

Bronchoalveolar lavage fluid (BALF) was obtained by intubating mouse tracheas through a small incision using a 20-gauge catheter. The bronchoalveolar space was washed by three sequential washes with 1 ml PBS containing 2 mM EDTA each. Cell-free BALF was obtained by centrifugation at 1500 rpm for 10 min at 4°C.

Mouse lungs were inflated by slowly instilling 1 ml 4% methanol-free PFA (Thermo Fischer Scientific) in PBS through the trachea. Whole lungs were dissected out in the inspiration phase and stored in 4% PFA overnight at 4^°^C in PBS before paraffin-embedding.

To isolate human PMBCs from Cohort I (healthy volunteers and AAV patients, used for cGAMP quantification), peripheral blood was collected into EDTA tubes (Sarstedt). PBMCs were isolated by Biocoll Separating Solution (Merck Millipore). Cells were washed twice in saline and counted. Cell pellets were immediately frozen at −80°C.

To isolate human PBMCs from Cohort II (healthy volunteers and AAV patients, used for gene set enrichment analysis), peripheral blood was collected into 4% sodium citrate. Within 15 min of collection, blood was diluted 1:2 with MACS rinsing buffer (PBS, 2 mM EDTA) and centrifuged on Histopaque 1077 at 900 g for 20 min at room temperature. Following centrifugation, PBMCs at the interface were removed, washed twice with MACS rinsing buffer, and then resuspended in 50 ml MACS running buffer (PBS, 2 mM EDTA, 0.5% BSA).

### Quantification of pulmonary hemorrhages

Hemoglobin (Hb) in the BALF was quantified spectrophotometry. 75 µl BALF was serially diluted in a 96-well plate and its optical density (OD) measured at 400 nm (Robles, Chowdhury, and Wax 2010) in a microplate reader with background subtraction at 670 nm. Values in the linear range below saturation (OD_400-670_ = 1.5) were selected and corrected for the dilution factor. Values were expressed as “corrected OD (Hb)”. Hb OD directly correlated to Hb concentration quantified by ELISA (Hemoglobin Assay Kit, Sigma Aldrich) (r^2^ = 0.9659, p = <0.0001).

### Neutrophil stimulation *in vitro*

Neutrophils were isolated from mouse bone marrow (femur and tibia) by Histopaque-based density gradient centrifugation as previously described (Swamydas et al. 2015). 2 x 10^5^ neutrophils in RPMI were seeded on poly-L-lysine-coated 12-mm glass coverslips. Neutrophils were incubated with 1 μM fMLP in RPMI or RPMI alone for 3 h at 37°C in a 5% CO_2_ humidified incubator. For the last 30 min, anti-MPO antibody (clones 6D1 and 6G4) was added at a final concentration of 5 µg/ml each. Cells were then carefully washed twice with warm RPMI and fixed in PBS containing 4% PFA for 30 min. Neutrophils were washed twice with PBS and stained with AlexaFluor 647-labelled goat anti-mouse IgG and DAPI in blocking solution (0.1M Tris solution pH 7.4 with 1% BSA, 1% gelatin from cold water fish skin and 0.3% Triton-X100). Cover slips were mounted on glass slides and images were acquired on a Zeiss LSM710 confocal microscope and analysed using FIJI-ImageJ software v. 2.0.0..

### Flow Cytometry

1-2×10^6^ cells were centrifuged at 1200 rpm for 5 min at 4°C. Cells were then resuspended in 100 µl PBS containing 3% FCS, 0.1% NaN_3_, 5% Octagam (human poly Ig) and fluorochrome-conjugated antibodies directed against different cell surface antigens for 20 min on ice in the dark. Antibodies are listed in Supplemental Table 4. Cells were washed and acquired in a LSR Fortessa flow cytometer (BD Biosciences). Sample acquisition was stopped in all different samples when an equal number of CaliBRITE beads (BD Biosciences) had been acquired. Dead cells and doublets were excluded with Hoechst33258 and standard procedures, respectively. Viable cells were then analyzed with FlowJo software v. 10.6.1.

Dimensionality reduction of flow cytometric data was performed using the Uniform Manifold Approximation and Projection for Dimension Reduction (UMAP) plug-in in FlowJo v. 10.6.1 and standard settings. Prior to UMAP, technical outliers were excluded using the FlowAI plug-in in FlowJo. Doublets and dead cells were excluded prior to UMAP. Specific cell populations were identified by standard gating as described in Fig. S2.

Reactive oxygen species (ROS) were quantified by flow cytometry. After staining for surface markers, cells were resuspended in 50 μl 0.5 mM dihydrorhodamine 123 (DHR123) in PBS for 30 min at 37°C and 5% CO_2_. Cells incubated with 1 μg/ml of PMA for 30 min at 37°C were used as positive controls. Samples not treated with DHR123 were used as negative controls.

### Giemsa and iron staining

Standard cytospins of BALF or FACS-sorted cells were stained with Giemsa to visualize cell morphology and RBC uptake. Slides were fixed in ice cold methanol for 10 mins at room temperature. Methanol was removed by washing slides in double distilled water (ddH_2_O). Afterwards, slides were stained with 20% Giemsa in ddH_2_O for 20 mins at RT. Slides were washed in ddH_2_O and mounted with ROTI Histokitt (ROTH). Images were taken using a standard light microscope.

Prussian blue stain (Iron Stain Kit, Abcam) was used to visualize iron in BALF cytospins according to manufacturer’s description. Slides were counterstained in nuclear fast red for 5 mins, washed in distilled water and mounted. Images were taken using a standard light microscope.

### cGAMP measurement in human PBMCs

cGAMP in human PBMCs was quantified with the 2’,3’-Cyclic GAMP ELISA Kit (Arbor assays) according to the manufacturer’s protocol. Briefly, 5 x 10^6^ thawed PBMCs were centrifuged and lysed in 50 µl sample diluent provided with the kit and assayed on a 96-well plate. For quantification of cGAMP in mouse samples, cell-free BALF supernatant from the first wash was centrifuged again at 6797 g for 15 min at 8°C. cGAMP was measured in undiluted BALF using the 2’,3’-Cyclic GAMP ELISA Kit (Cayman Chemical) according to the manufacturer’s protocol. Sample concentrations were calculated with GraphPad Prism 8 using 4PL regression to interpolate the standard curve.

### Histopathological analysis and lung injury score

Histopathological quantification was performed by a qualified pathologist (P.B.) in a blinded manner on 1 µm thick microtome sections of paraffin-embedded whole-lungs stained with H&E and scanned at a 40x magnification using a bright-field scanner (Aperio AT2, Leica). Morphological changes and immune cell infiltration in the lungs were graded as previously described (Li et al. 2014). In brief, the lung injury score was composed of 3 parameters: immune cell infiltration, tissue destruction and the number of lobes affected. Scores were: 0 (no inflammation), normal lung structure; 1 (slight), very few and only focal inflammatory cells, no or minimal tissue destruction; 2 (medium), larger areas also with tissue destruction; 3 (strong), large areas with inflammatory infiltrates and tissue destruction, up to one lobe involved; 4 (extensive), as 3 but involving more than 1 whole lobe.

### Confocal microscopy of the lung

To investigate endothelial cell death, we injected 5 µg AF647-conjugated anti-CD31 antibody i.v. 5 mins prior to humane killing of mice. Lungs were inflated with 2% low-melting agarose solution at 37°C. Lungs were dissected out, cooled for 10 minutes at 4°C and finally embedded in 4% agarose. Lung sections of 150 µm thickness were cut using a vibrating-blade microtome. Sections were mounted in PBS containing 1 µg/ml propidium iodide and immediately analyzed on a LSM 710 Zeiss confocal microscope. Five z-stacks covering 15 µm were taken per image. The first photograph was taken at 25 µm from the surface to avoid analysing cells damaged during the cutting process. Data was analyzed using FIJI-ImageJ software v. 2.0.0.

### Optical clearing of the lung and light-sheet microscopy

Three hours after AAPV induction, mice were injected intravenously (i.v.) with 50 µg Texas red-labeled albumin in 100 µl PBS. One hour later, animals were killed in CO_2_ / air (50/50 (v/v)) and exsanguinated by an incision in the right femoral artery. Mice were perfused in PBS though the left ventricle at a flow speed of 10 ml/min for 90 sec. Lungs were subsequently explanted and fixed in 4% methanol-free PFA overnight at 4°C.

Optical clearing was achieved as previously described (Klingberg et al. 2017) by transfer of the samples into 99% ethanol for 3 h at room temperature. Samples were then transferred into ethyl cinnamate and incubated for further 5 h. Image stacks of the cleared lungs were acquired with a step size of 20 µm using a custom build light-sheet-fluorescence-microscope (LaVision Biotec). Samples were excited with two light-sheets at 561 nm and recorded using an emission filter at 620/60 nm to detect albumin depositions. Texas red-labeled albumin deposition was quantified in FIJI-Image J by physician-aided thresholding, resulting in the scale bar within the picture. Thresholds were set to equal values for all samples. Files were exported in nrrd format and imported into ParaView v5.7.0 for 3D-reconstruction.

### Extracellular DNA quantification in BALF

DNA in 100 µl of undiluted cell-free BALF (first wash) was quantified in DNase-free 96-well plates (TPP^®^) using Quant-iT^TM^ Pico Green^TM^ dsDNA Assay Kit (Thermo Fisher Scientific) following manufacturers’ instructions.

### Extracellular DNA purification and analysis

DNA in cell-free BALF was purified using standard phenol-chloroform-isoamyl procedures, precipitated in 3 M sodium acetate and isopropanol at −80°C for at least 1 h, and finally resuspended in DNase free water. DNA concentration was measured using a spectrophotometer.

Recombinant 6xHis-tagged murine Trex1 was expressed in *E. coli* Rosetta DE3 and purified by Ni-NTA affinity chromatography as previously described (Gehrke et al. 2013). For digestion experiments, 1 µg of purified DNA was incubated with 40ng recombinant murine Trex1, 1 U DNase I or without enzyme in 25 µl of digestion buffer (20 mM Tris pH 7.5, 10 mM MgCl_2_, 60 mM KCl). Digestion was performed at 37 °C for 30 min followed by enzyme inactivation at 75 °C for 10 min. Half of the digested samples were run on a 1.5 % agarose gel at 110 V for 1h 30min. The gel was stained with SYBR Gold (Thermo Fisher Scientific) and imaged on a ChemiDoc XRS+(Bio-Rad). Trex1-resistant (but DNase I-sensitive) modified 60bp DNA was used as controls as previously described (Gehrke et al. 2013).

The ratio of mitochondrial to nuclear extracellular DNA purified from cell-free BALF was determined by qRT-PCR as previously described (Quiros et al. 2017). qRT-PCR was performed on the purified DNA with the iTaq Universal SYBR® Green Supermix (Bio-Rad) on a QuantStudio 6 Flex System (Thermo Fisher Scientific). Primer sequences for the mitochondrial genes *Nd1* and *16s*, and the nuclear gene *Hk2* are listed in Supplement table 3. Relative gene abundance was calculated by the ^ΔΔ^Ct method.

### Analysis of IFN type-I signature in mice

Analysis of ISGs was performed as previously described (Luksch et al. 2019). Total RNA from mouse lungs was extracted using RNeasy Mini Kit (Qiagen, Hilden, Germany) according to the manufacturer’s instructions. RNA concentration and purity were determined spectrophotometrically. 2 µg RNA was reverse transcribed with Moloney murine leukemia virus reverse transcriptase, RNasin and dNTPs (all Promega, Mannheim, Germany) in 35 µl reaction according manufactures instructions. Quantitative RT-PCR was performed using Quantstudio 5 (Thermo Fisher Scientific, Berlin, Germany) and GoTag^®^qPCR Master Mix with SYBR green fluorescence (Promega, Mannheim, Germany). PCR primer sequences were retrieved from online Primer Bank database (Spandidos et al., 2009). Expression of genes was normalized with respect to each housekeeping gene using the ^ΔΔ^Ct method for comparing relative expression (up-regulation > 2.0 and down-regulation < 0.5).

### Gene set enrichment analysis (GSEA)

Gene set enrichment analysis was performed as previously described (Lyons et al. 2010). Briefly, RNA was extracted from 5×10^6^ human PBMCs from Cohort II using AllPrep RNEasy kits (Qiagen) and the small RNA-containing column flow-through collected. For microarray analyses, 200 ng aliquots of total RNA were labelled and hybridized to Affymetrix Gene ST 1.1 expression arrays according to the manufacturer’s instructions. Microarray data was stored as CEL files and read into R. Raw data was pre-processed using the BioConductor suite of packages. The oligo package was used to normalize the data and create an expression set. Raw values were log transformed and normalized using robust multiarray averaging (RMA). The arrayQualityMetrics package was used to assess outliers. Probes with no annotation (Entrez gene ID) were removed. Where multiple probes mapped to a common gene, the probe with the largest variance was analyzed. ComBat (SVA package) was used to correct for batch affects. GSEA was performed to assess for enrichment in the interferon pathway. The package ComplexHeatmap was used to generate the heatmap of leading-edge genes using Z normalized data.

### cGAS activation assay

Radiolabeled cGAS activity assays were performed as previously described (Civril et al. 2013). Briefly, 80 ng of DNA was mixed with 25 µM ATP, 25 µM GTP and trace amounts of [⍺^32^P] ATP (Hartman Analytic) in reaction buffer (50 mM HEPES pH 7.5, 5 mM MgCl_2_, 10 mM NaCl, 1 mM DTT). Reactions were started by addition of 1 µM mouse cGAS and incubated for 45 min at 37°C. Reaction products were spotted on thin layer chromatography (PEI-Cellulose F plates, Merck) and 1 M (NH_4_)_2_SO_4_/1.5 M KH_2_PO_4_ pH 3.8 was used as running buffer. TLC plates were analyzed by phosphor imaging (Typhoon FLA 9000, GE Healthcare). Genomic DNA (gDNA) and 60bp synthetic DNA (dsDNS) were used as controls.

### *Ex vivo* bioluminescence imaging and quantification

Lungs of *Ifnβ*-Luc reporter mice were taken after 3 days of vasculitis induction and digested using DNase I (50 U/ml) and collagenase IV (1 mg/ml) for 30 mins at 37°C. Afterwards, red blood cells were lysed by ACK (Ammonium-Chloride-Potassium) buffer for 10 mins at room temperature. Cells were washed, surface stained and sorted by FACS. Cells numbers were then adjusted and cells were plated in a white 96-well plate. 20 µl of passive lysis buffer (Promega) was added per well and plates shaked for 5 mins at 1400 rpm on a rocking plate. Luminescence was quantified using a plate reader for 1000 ms before and after adding 100 µl of Luciferin substrate assay to each well.

### Quantification and statistical Analysis

Processing of data for GSEA has been described in section “Gene set enrichment analysis (GSEA)”. All other raw data was processed with Excel (version 16.26) and GraphPad Prism (version 8.1.2). Experiments were performed at least two times with a minimum of ≥5 animals per group. Results are presented as mean ± SEM and no individual data points were excluded under any circumstances. Student’s *t-*test (with Welch’s correction for non-parametric data) was used for two-group comparisons, and one- or two-way ANOVA with Tukey post-test for multiple group comparison (Kruskal-Wallis test for non-parametric data). Statistics are stated as: *, p < 0.05; **, p < 0.01; ***, p < 0.001.

### Online supplemental material

Fig. S1 shows validation of MPO-specific IgGs by ELISA, and additional immunological features in the lung of mice undergoing AAPV. Figure S2 shows the flow cytometric gates for cell identity. Figure S3 shows additional ISG in the lungs of WT and STING^−/−^ mice undergoing AAPV. Figure S4 shows flow cytometric gating strategy and phenotype of different myeloid cells and macrophages, as well as iron staining in macrophages and neutrophils.

Table S1 and S2 show demographic and clinical characteristics of patient cohorts I and II, respectively. Table S3 shows primer sequences. Table S4 shows antibodies used for flow cytometry.

## Abbreviations

AAPV: ANCA-associated pulmonary vasculitis
AAV: ANCA-associated vasculitis
Alv.: alveolar
ANCA: Anti-neutrophil cytoplasmic antibody
BALF: broncho alveolar lavage fluid
GSEA: gene set enrichment analysis
Hb: Hemoglobin
infl.: inflammatory
i.t.: intratracheal
i.p.: intraperitoneal
MPO: Myeloperoxidase
PR3: Proteinase 3
SAVI: STING-associated vasculopathy with onset in infancy

## Acknowledgments

We thank Christine Schmidt, Daniela Klaus, Saskia Schmitz, Anita van Esch, and the staff at the Flow Cytometry Core Facility of the Faculty of Medicine, University of Bonn, for excellent technical assistance. Work in the laboratories of N.G., C.K., L.T., A.R.-W., K.-P.H., G.H., and E.B. was funded by the Deutsche Forschungsgemeinschaft (DFG, German Research Foundation) – Grant Number 369799452 –. Work in the laboratories of N.G., C.K., L.T., G.H., and E.B. is funded by the Deutsche Forschungsgemeinschaft (DFG, German Research Foundation) under Germany’s Excellence Strategy – EXC 2151 – 390873048. C.K., P.L., P.H., D.J received funding from the European Union’s Horizon 2020 research and innovation program (grant agreement no. 668036 “RELENT”). PB was supported by the German Research Foundation (DFG, Project IDs 322900939, 454024652, 432698239 & 445703531), European Research Council (ERC Consolidator Grant No 101001791), and the Federal Ministry of Education and Research (BMBF, STOP-FSGS-01GM1901A). C.C.O.M. is supported as a Cancer Research Institute/Eugene V. Weissman Fellow.

## Author contributions

N.K., S.F.V, C.K., K.M., K.M.K, H.L, A.M.C.B., C.C.O.M., S.A.I.W. and T.Z. planned and performed experiments, and analyzed data. S.O. and P.B. performed and analyzed histopathology. P.H. and D.H. generated the monoclonal antibodies against murine MPO and contributed to the development of the mouse vasculitis model. L.T. organized patient cohort 1 used for cGAMP measurements. P.K. and P.A.L. organized and analyzed patient cohort 2 used to identify IFN-I signature. N.G., C.K., D.E.J., A.A, E.B, G.H., K-P.H., A.R.-W. and L.T. contributed to planning of experiments and supervised work in their laboratories. N.G. and C.K. coordinated the project. N.K., S.F.V. and N.G. wrote the manuscript with contributions from all authors.

