## Supplementary Figures for "Monocyte-derived macrophages aggravate pulmonary vasculitis via cGAS/STING/IFN-mediated nucleic acid sensing"

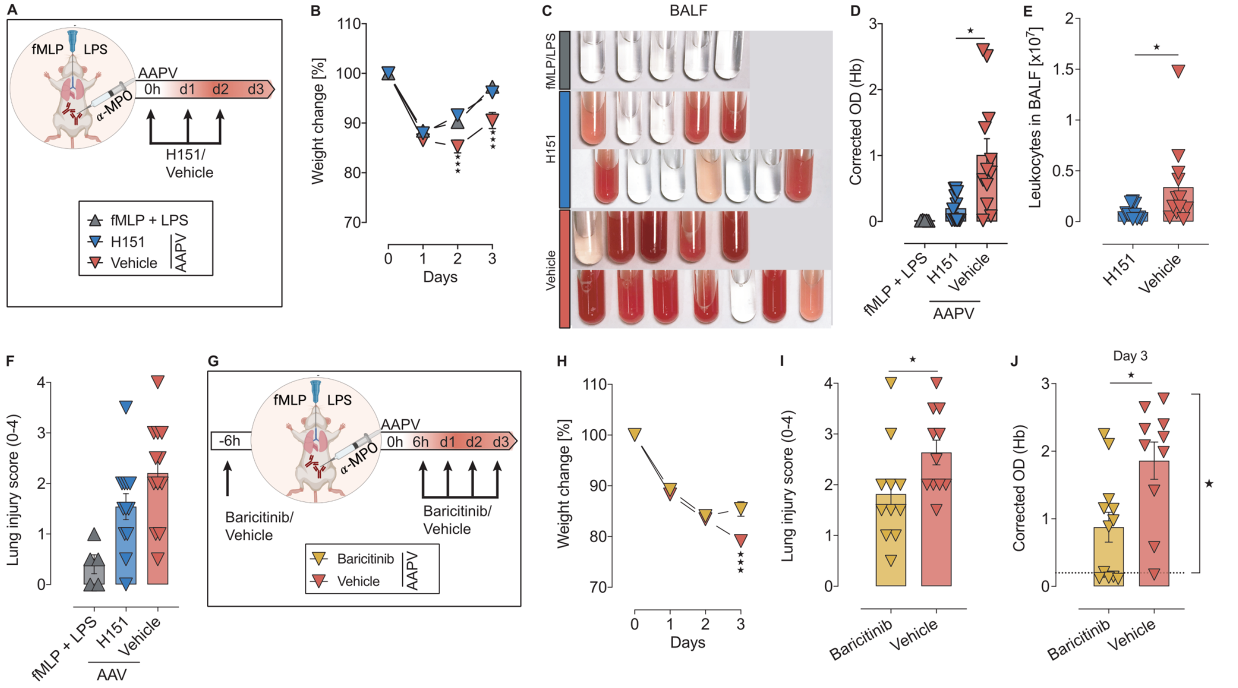


Figure 7. **Pharmacological inhibition of STING or JAK/STAT pathways ameliorates AAPV in mice. (A)** Schematic representation of experiment with H151 STING inhibitor. **(B-D)** Weight loss kinetics (B), hemorrhages in BALF (C, D), leukocyte infiltration (E) and lung injury score (F) in mice treated as in (A). **(G)** Schematic representation of experiment with JAK1/2 inhibitor baricitinib. **(H-J)** Weight loss (H), lung injury score (I) and hemorrhages in the BALF (J) of WT mice treated as in (A). Each dot is an individual mouse (D-F, I, J). Bar and line graphs show mean ± SEM. Data is pooled from 2 independent experiments. *, P < 0.05; **, P < 0.01.***, P < 0.001. Hb, hemoglobin.

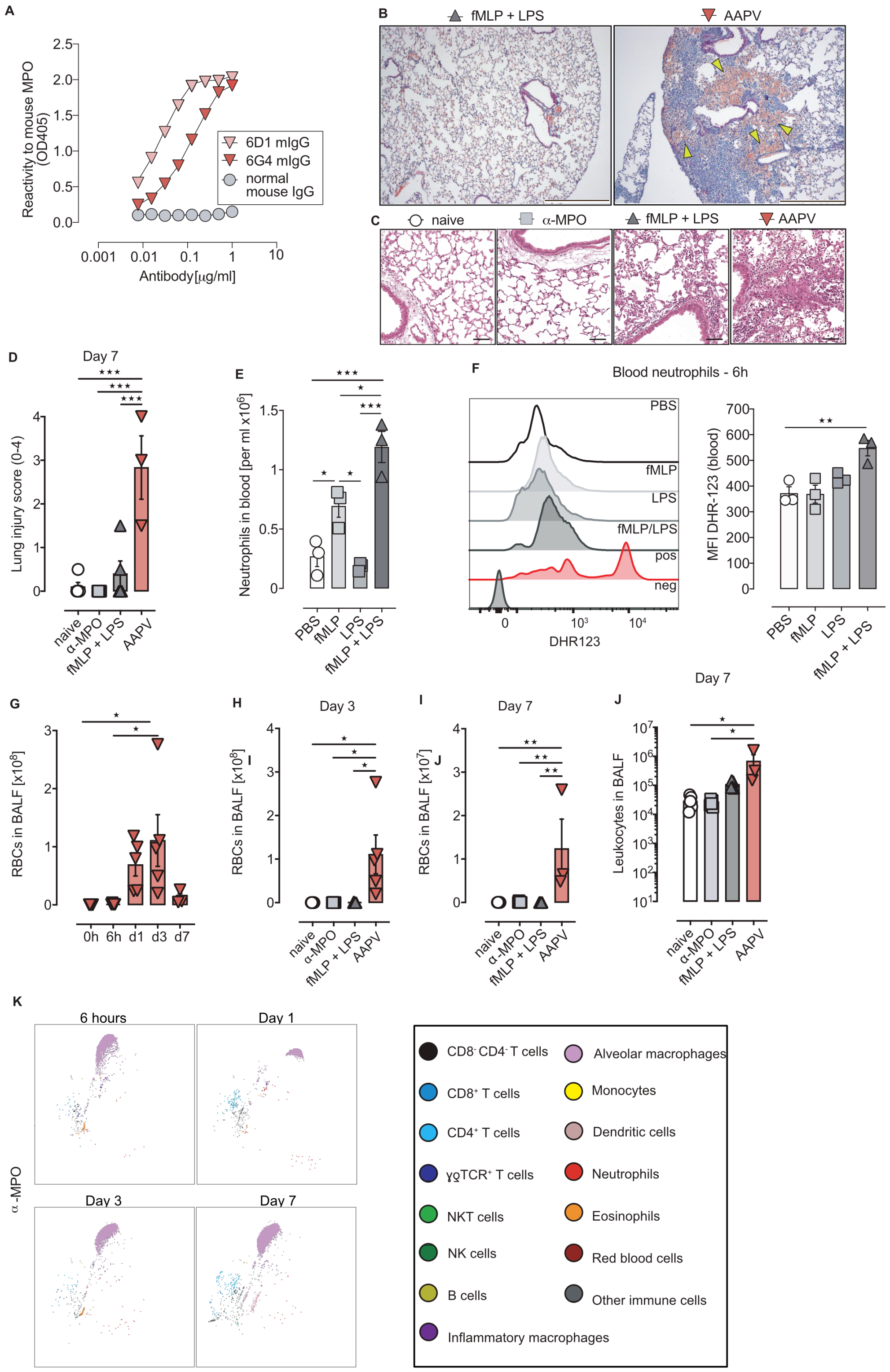


Figure S1. **Anti-MPO antibodies and low dose bacterial ligands synergize to induce severe pulmonary vasculitis in a novel mouse model of pulmonary vasculitis. (A)** ELISA for recombinant MPO with the mAb clones 6D1 and 6G4 indicating specificity for mouse MPO. **(B, C)** H&E stainings of lung cryosections of mice treated as in Fig. 2A. Yellow arrow heads indicate areas with hemorrhages. **(D)** Lung injury score 7 days after treatment of mice as in Fig. 2A. **(E)** Quantitative flow cytometry analysis of circulating neutrophils in blood of WT mice 6 h after indicated treatments. **(F)** Representative flow cytometric histograms (left) and quantification of ROS production (DHR123 MFI) by blood neutrophils from mice treated as in (E). pos, positive control (PMA); neg, fluorescence minus one (FMO) for DHR123 in untreated neutrophils. **(G-J)** RBC (G, H, I) and leukocyte (J) counts in BALF of the indicated mice. **(K)** UMAP dimensionality reduction of immune cells in BALF of mice treated only with anti-MPO antibodies using identity gates in Fig. S2. Results for individual mice are shown as dots. Data is representative of at least 2 independent experiments (mean ± SEM). *, $P < 0.05$; **, $P < 0.01$; ***, $P < 0.001$. RBCs, red blood cells.

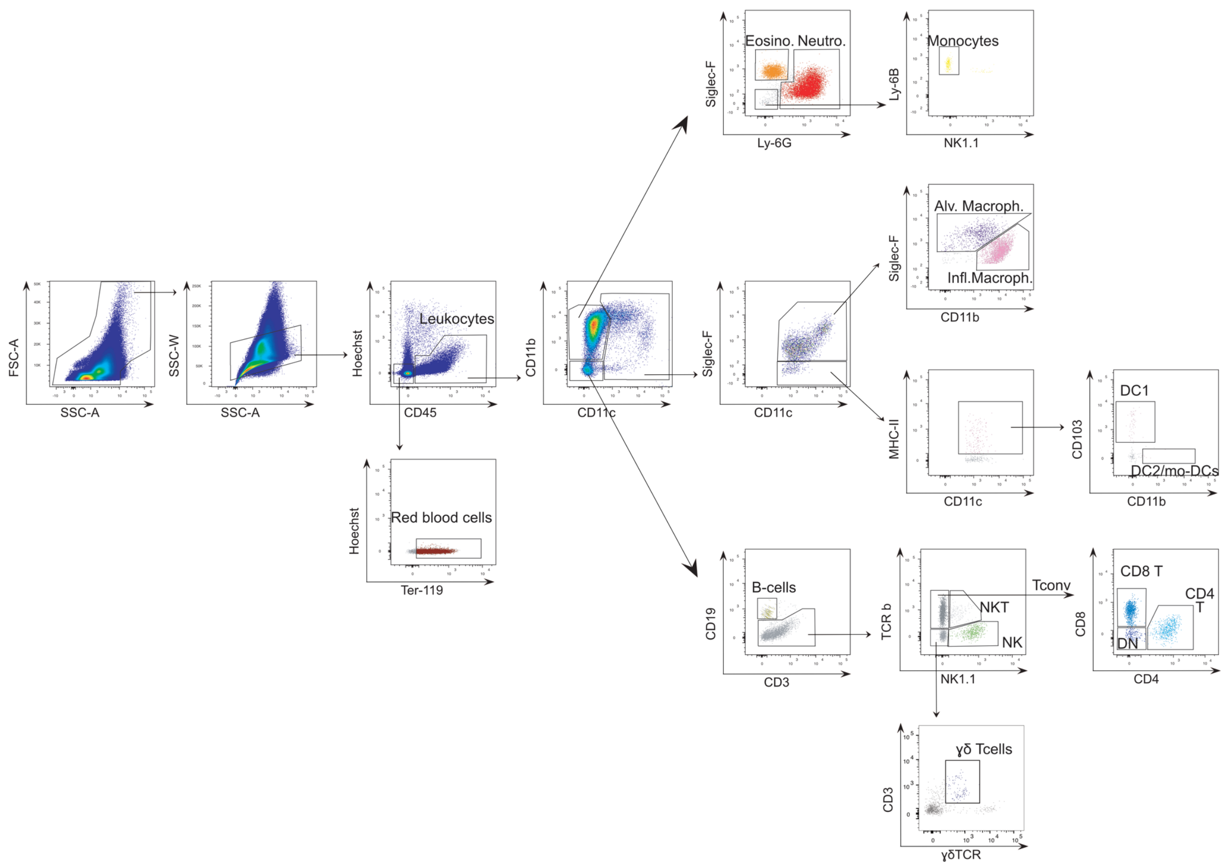


Figure S2. **Cell identity gates used in this study.** Hierarchy of flow cytometry dot plots and gates of lung singe-cell suspensions from wild-type mice 3 days after AAPV induction used to identify different cell populations.

| | |
|---|---|
| Leukocytes: | single Hoechst33258$^{-}$ CD45$^{+}$ events. |
| Red blood cells: | single Hoechst33258$^{-}$ CD45$^{-}$ Ter119$^{+}$ events. |
| Eosinophils: | CD11b+ CD11c$^{-}$ Siglec-F$^{+}$ Ly-6G$^{-}$ leukocytes. |
| Neutrophils: | CD11b+ CD11c$^{-}$ Siglec-F$^{-/+}$ Ly-6G$^{+}$ leukocytes. |
| Monocytes: | CD11b+ CD11c$^{-}$ Siglec-F$^{-}$ Ly-6G$^{-}$ NK1.1$^{-}$ Ly-6B$^{+}$ leukocytes. |
| Alveolar Møs: | CD11c$^{+}$ Siglec-F$^{hi}$ CD11b$^{-/int}$ leukocytes. |
| Inflammatory Møs: | CD11c$^{+}$ Siglec-F$^{int}$ CD11b$^{int/hi}$ leukocytes. |
| cDC1: | Siglec-F$^{-}$ CD11c$^{+}$ MHC-II$^{+}$ CD103$^{+}$ CD11b$^{-}$ leukocytes. |
| cDC2/mo-DCs: | Siglec-F$^{-}$ CD11c$^{+}$ MHC-II$^{+}$ CD103$^{-}$ CD11b$^{+}$ leukocytes. |
| B cells: | CD11c$^{-}$ CD11b$^{-}$ CD3ε$^{-}$ CD19$^{+}$ leukocytes. |
| CD4 α/β T cells: | CD11c$^{-}$ CD11b$^{-}$ CD3ε$^{+}$ CD19$^{-}$ TCRβ$^{+}$ NK1.1$^{-}$ CD8$^{-}$ CD4$^{+}$ leukocytes. |
| CD8 α/β T cells: | CD11c$^{-}$ CD11b$^{-}$ CD3ε$^{+}$ CD19$^{-}$ TCRβ$^{+}$ NK1.1$^{-}$ CD8$^{+}$ CD4$^{-}$ leukocytes. |
| DN α/β T cells: | CD11c$^{-}$ CD11b$^{-}$ CD3ε$^{+}$ CD19$^{-}$ TCRβ$^{+}$ NK1.1$^{-}$ CD8$^{-}$ CD4$^{-}$ leukocytes. |
| γδ T cells: | CD11c$^{-}$ CD11b$^{-}$ CD3ε$^{+}$ CD19$^{-}$ TCRβ$^{-}$ γδ TCR$^{+}$ |
| NK cells: | CD11c$^{-}$ CD11b$^{-/+}$ CD3ε$^{+}$ CD19$^{-}$ TCRβ$^{+}$ NK1.1$^{+}$ leukocytes. |
| NKT cells: | CD11c$^{-}$ CD11b$^{-}$ CD3ε$^{+}$ CD19$^{-}$ TCRβ$^{+}$ NK1.1$^{+}$ leukocytes. |

Kessler at al.  Figure S3

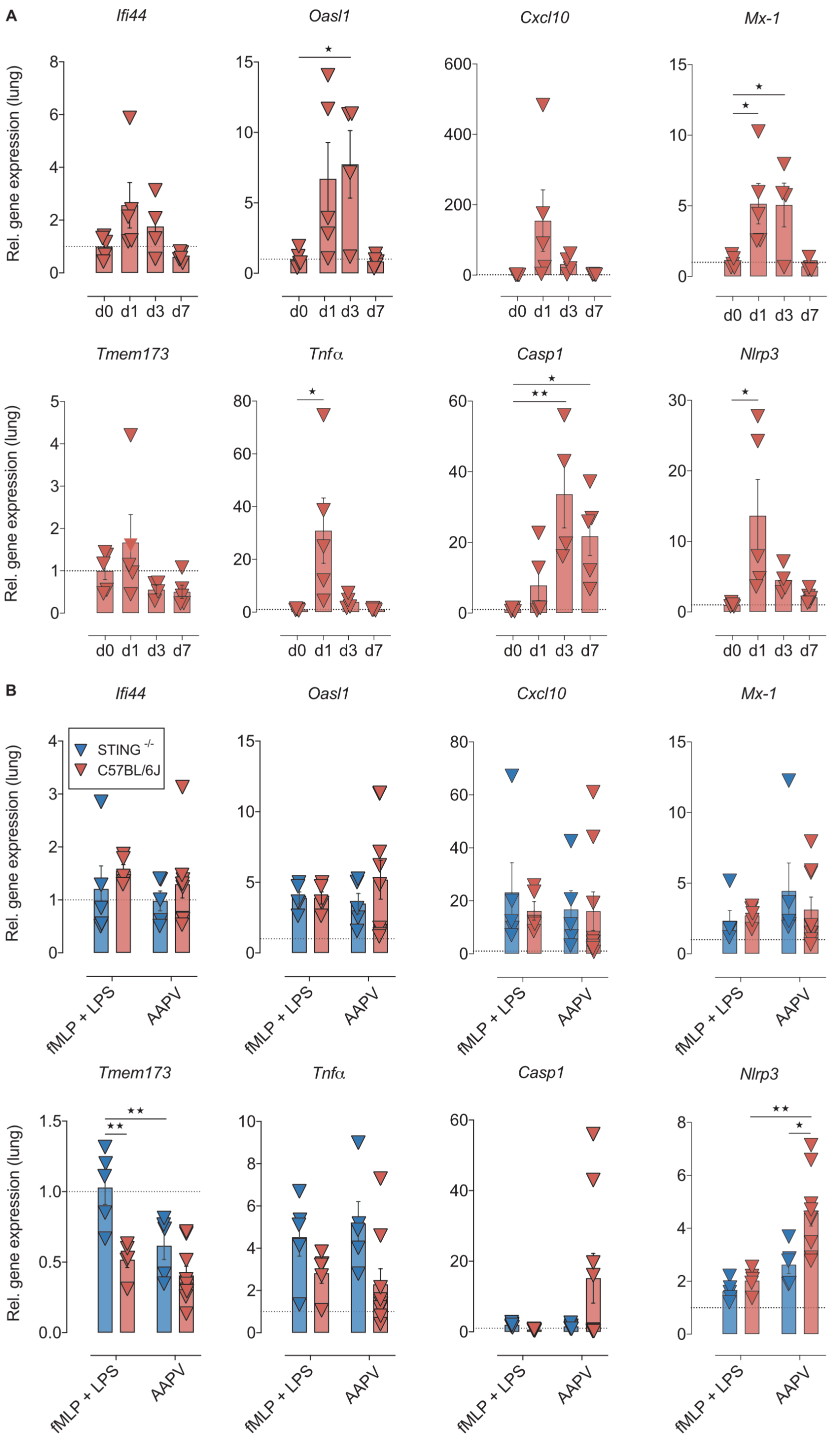

Figure S3. **IFN-I signature in the lung of mice with autoimmune AAPV. (A)** Relative gene expression in the lungs of C57BL/6J at the indicated time points after AAPV induction. **(B)** Relative gene expression in the lungs of the indicated mice 3 days after treatment as indicated.
Each dot is a single mouse (n = 5-8 mice/group). Bars represent mean ± SEM. *, $P < 0.05$; **, $P < 0.01$.

Kessler at al. Figure S4

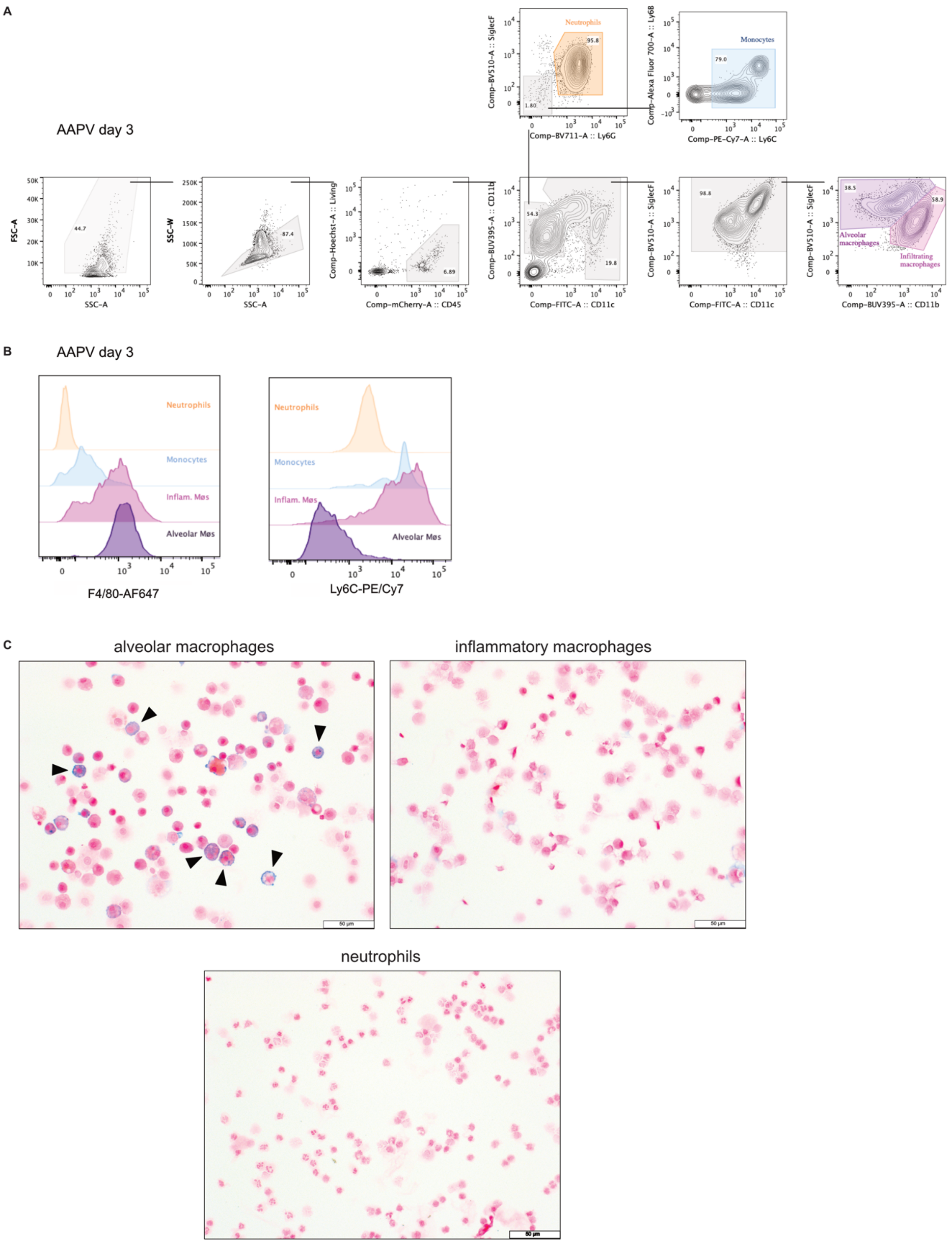


Figure S4. **Characteristics of tissue-resident alveolar macrophages and infiltrating inflammatory macrophages in the lung of mice with autoimmune vasculitis. (A, B)** Flow cytometric gating strategy, F4/80 and Ly6C expression of the indicated macrophage and myeloid cell populations. **(C)** Iron staining or FACS-sorted alveolar macrophages, infiltrating macrophages, and neutrophils as in (A). Arrowheads mark examples of Alv. Møs with iron staining.

Table S1. **Patient cohort I. Demographic and clinical characteristic of AAPV patients (related to Fig. 1).**

| Characteristics | AAV patients (n = 10) |
|---|---|
| Age - years mean ± SD | 60 ± 13 |
| Sex | |
| Female - no (%) | 3 (30) |
| Male - no (%) | 7 (70) |
| Diagnosis | |
| GPA - no (%) | 8 (80) |
| EGPA - no (%) | 1 (10) |
| MPA - no (%) | 1 (10) |
| cANCA positive at baseline - no (%) | 6 (60) |
| pANCA positive at baseline - no (%) | 1 (10) |
| Therapy at baseline | |
| Glucocorticoids - no (%) | 5 (50) |
| Rituximab - no (%) | 4 (40) |
| Methotrexate - no (%) | 4 (40) |
| Mycophenolate mofetil - no (%) | 1 (10) |
| Azathioprine - no (%) | 1 (10) |
| Clinical manifestation | |
| Ear, nose, throat - no (%) | 4 (40) |
| Pulmonary - no (%) | 4 (40) |
| Renal - no (%) | 3 (30) |
| Cutaneous - no (%) | 3 (30) |
| Musculoskeletal - no (%) | 4 (40) |
| Neurologic - no (%) | 5 (50) |
| Digital ischemia - no (%) | 2 (20) |
| Ophthalmic | 2 (20) |
| Duration since diagnosis - years mean ± SD | 9.0 ± 7.6 |

| | |
|---|---|
| Laboratory results at baseline | |
| C-reactive protein - mg/l mean ± SD | 31.5 ± 87.7 |
| Creatinine - mg/l mean ± SD | 0.87 ± 0.45 |
| eGFR - ml/min/1.73m$^2$ mean ± SD | 65.4 ± 13.7 |

no, number

Table S2. **Patient cohort II. Demographic and clinical characteristic of AAPV patients**
(Related to Fig. 1)

| **Characteristics** | **MPO- AAV patients (n = 21)** | **Healthy Controls (n= 47)** |
|---|---|---|
| Age - years mean ± SD | 64 ± 14 | 47 ± 15 |
| Sex | | |
| Female - no (%) | 13 (62) | 28 (60) |
| Male - no (%) | 8 (38) | 19 (40) |
| Diagnosis | | |
| GPA - no (%) | 5 (24) | |
| EGPA - no (%) | 4 (19) | |
| MPA - no (%) | 12 (57) | |
| Therapy at baseline | | |
| Adalimumab - no (%) | 3 (14) | |
| Mycophenolate mofetil - no (%) | 3 (14) | |
| Cyclophosphamide - no (%) | 12 (57) | |
| Rituximab - no (%) | 1(5) | |
| Clinical manifestation | | |
| Ear, nose, throat - no (%) | 6 (29) | |
| Pulmonary - no (%) | 6 (29) | |
| Renal - no (%) | 11 (55) | |
| Laboratory results at baseline | | |
| C-reactive protein - mg/l mean ± SD | 53 ± 58 | |
| Creatinine - mg/l mean ± SD | 0.29 ± 0.33 | |

Table S3. **Primer sequences (*M. musculus* genes)**
(Related to methods)

| Gene | Forward | Reverse |
|---|---|---|
| *16s* | 5´-CCG CAA GGG AAA GAT GAA AGA C-3´ | 5´-TCG TTT GGT TTC GGG GTT TC 3´ |
| *Casp1* | 5'-GCT GCC TGC CCA GAG CAC AAG-3' | 5'-CTC TTC AGA GTC TCT TAC TG-3' |
| *Mb21d1* | 5'-GAG GCG CGG AAA GTC GTA A-3' | 5'-TTG TCC GGT TCC TTC CTG GA-3' |
| *Cxcl10* | 5'-CCA AGT GCT GCC GTC ATT TTC-3' | 5'-GGC TCG CAG GGA TGA TTT CAA-3' |
| *Eef2* | 5'-CCG ACT CCC TTG TGT GCA A-3' | 5'-AGT TCA GGT CGT TCT CAG AGA G -3' |
| *Hk2* | 5´-GCC AGC CTC TCC TGA TTT TAG TGT-3´ | 5´-GGG AAC ACA AAA GAC CTC TTC TGG-3´ |
| *Hprt1* | 5'-TCA GTC AAC GGG GGA AT AAA-3' | 5'-GGG GCT GTA CTG CTT AAC CAG-3' |
| *Ifi44* | 5' –AAC TGA CTG CTC GCA ATA ATG T-3' | 5'-GTA ACA CAG CAA TGC CTC TTG T-3' |
| *Ifit2* | 5'-GCT CTG GAA AAG GAC CCG AA-3' | 5'-GCT TCA GTG CCA AGA GGA CT-3' |
| *Ifnβ* | 5'-TCC GAG CAG AGA TCT TCA GGA A-3' | 5'-TGC AAC CAC CAC TCA TTC TGA G-3' |
| *Il1β* | 5'-GAA ATG CCA CCT TTT GAC AGT G-3' | 5'-TGG ATG CTC TCA TCA GGA CAG-3' |
| *Il-6* | 5'-CTG CAA GAG ACT TCC ATC CAG-3' | 5'-AGT GGT ATA GAC AGG TCT GTT GG-3' |
| *Isg15* | 5'-CAA TGG CCT GGG ACC TAA AG-3' | 5'-CTG TAC CAC TAG CAT CAC TGT G-3' |
| *Mx1* | 5'-AAC CCT GCT ACC TTT CAA-3' | 5'-AAG CAT CGT TTT CTC TAT TTC-3' |
| *Nd1* | 5´-CTA GCA GAA ACA AAC CGG GC-3´ | 5´-CCG GCT GCG TAT TCT ACG TT-3´ |
| *Nlrp3* | 5'-ATC AAC AGG CGA GAC CTC TG-3' | 5'-GTC CTC CTG GCA TAC CAT AGA-3' |
| *Oasl1* | 5'-AGC TCC GAG GTC TAC GCA A-3' | 5'-GGG GCA CTT GTC TCT CAC AT-3' |
| *Rpl13a* | 5'-AGC CTA CCA GAA AGT TTG CTT AC-3' | 5'-GCT TCT TCT TCC GAT AGT GCA TC-3' |
| *Tmem173* | 5'-CTG CTG ACA TAT ACC TCA GTT G-3' | 5'-GAG CAT GTT GTT ATG TAG CTG-3' |
| *Tnfα* | 5'-CCT GTA GCC CAC GTC GTA G-3' | 5'-GGG AGT AGA CAA GGT ACA ACC C-3' |

Table S4. **Antibodies used for flow cytometry against mouse antigens**
(Related to methods)

| ANTIBODY | SOURCE | IDENTIFIER |
|---|---|---|
| anti-CD45 AF594 (clone 30-F11) | BioLegend, San Diego, USA | Cat# 103144; RRID:AB_2563458 |
| anti-NK1.1 AF647 (clone PK136) | BioLegend, San Diego, USA | Cat# 108720; RRID:AB_2132713 |
| anti-Ly6G AF711 (clone 1A8 | BioLegend, San Diego, USA | 127643; RRID:AB_2565971 |
| anti-TCRβ APC/eF780 (clone H57-597) | eBioscience, San Diego, CA, USA | Cat# 47-5961-82; RRID:AB_1272173 |
| anti-CD11b BUV395 (clone M1/70) | Becton, Dickinson and Co., NJ, USA | Cat# 563553; RRID:AB_2738276 |
| anti-CD4 BUV737 (clone GK1.5) | Becton, Dickinson and Co., NJ, USA | Cat# 564298; RRID:AB_2738734 |
| anti-CD8a BUV805 (clone 53-6.7) | Becton, Dickinson and Co., NJ, USA | Cat# 564920; RRID:AB_2716856 |
| anti-SiglecF BV510 (clone E-50-2440) | Becton, Dickinson and Co., NJ, USA | Cat# 740158; RRID:AB_2739911 |
| anti-CD44 BV605 (clone IM7) | BioLegend, San Diego, USA | Cat# 103047; RRID:AB_2562451 |
| anti-CD3e BV650 (clone 17A2) | BioLegend, San Diego, USA | Cat# 100229; RRID:AB_11204249 |
| anti-Ly6B2 AF700 (clone 7/4) | Bio-Rad, Hercules, CA, USA | Cat# MCA771A700T; RRID:AB_1102792 |
| anti-CD19 BV786 (clone 6D5) | BioLegend, San Diego, USA | Cat# 115543; RRID:AB_11218994 |
| anti-MHCII FITC (clone M5/114.15.2) | BioLegend, San Diego, USA | Cat# 107606; RRID:AB_313321 |
| anti-CD103 BV421/PB (clone 2E7) | BioLegend, San Diego, USA | Cat# 121418; RRID:AB_2128619 |
| anti-TCRɣδ PE, (clone GL3) | Becton, Dickinson and Co., NJ, USA | Cat# 553178; RRID:AB_394689 |
| anti-CD11c Pe Cy7 (clone N418) | eBioscience, San Diego, CA, USA | Cat# 25-0114-82; RRID:AB_469590 |
| anti-Ter119 PerCPCy5.5 (clone Ter119) | BioLegend, San Diego, USA | Cat# 116228; RRID:AB_893636 |
